# BCL11B targeting in tumor CD8^+^ T cells amplifies anti-tumor response by blocking exhaustion while promoting stemness and cytotoxicity

**DOI:** 10.64898/2026.08.03.742578

**Authors:** Leonardo Silvane, Tomas Zelenka, Divya P. Talada, Valeriu B. Cismasiu, Shamima Islam, Raghwendra P. Singh, Zefanias Ngove, Sayan Chakraborty, MacLean S. Hall, Jamie L. Blauvelt, Erika Eksioglu, Soraya Zorro Manrique, Joseph O. Johnson, Alyssa N. Obermayer, Alex Alfaro, Wen Huang, Amod Sarnaik, Ahmad A. Tarhini, John E. Mullinax, Erin George, Patrick Hwu, Eduardo Davila, Jose R. Conejo-Garcia, Yenan T. Bryceson, Dung-Tsa Chen, Timothy I. Shaw, Shari Pilon-Thomas, Dorina Avram

## Abstract

Tumor infiltrating CD8^+^ T cells (TILs) progress to a state of terminal exhaustion (Ttex) which have impaired functionality and are nonrenewable. However their precursors (Tpex) are renewable and can generate efficient effector cells. We started from the observation that melanoma patients undergoing therapy with checkpoint inhibitors show increased survival when their T cells have low *BCL11B* mRNA. In line with this, ablation of *Bcl11b* in CD8^+^ TILs conferred a superior anti-tumor response in murine melanoma and ovarian cancer models. *Bcl11b* KO TILs failed to progress to the Ttex state and retained elevated stemness. Bcl11b exerted its role by repressing expression of essential transcription factors (TF) controlling stemness, and conversely by promoting expression of exhaustion-associated TFs and inhibitory receptor genes, through complex epigenetic control. In addition, *Bcl11b* KO CD8^+^ T cells showed increased Ag-specific cytolytic activity and elevated Gzmb and Prf1 proteins, but no increase in their mRNAs, however presented higher expression of genes with role in translation. Furthermore, CRISPR-CAS9-mediated deletion of BCL11B in human TILs from a patient with poor response to adoptive cell therapy with autologous TILs, improved their cytolytic activity and promoted expression of the stemness-associated TF TCF1, underlying its potential therapeutic use.

**HIGHLIGHTS:**

– Adoptive transfer of *Bcl11b* KO CD8^+^ TILs surpasses WT in tumor burden reduction
– *Bcl11b* ablation reprograms TILs and impairs the progression to Ttex state
– *Bcl11b* KO CD8^+^ T cells have elevated cytotoxicity and kill only Ag-MHCI targets
– *BCL11B* deletion in nonresponder ACT-TIL improves cytolytic activity and elevates TCF1

**GRAPHICAL ABSTRACT:** 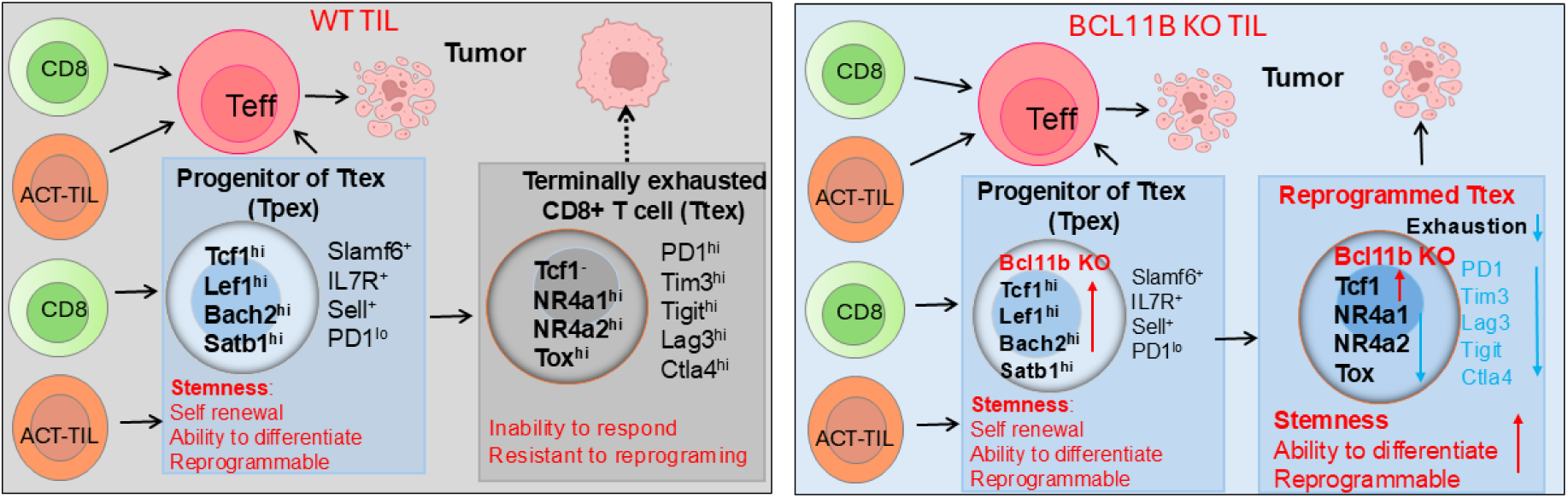

## Introduction

Immunotherapy approaches have revolutionized cancer treatment and have improved patient survival. Initial success was achieved by immune check point blockade (ICB) against the co-inhibitory receptors PD-1 and CTLA-4, with around 40% success rate in melanoma^1^. Furthermore, adoptive cell therapies (ACT) with autologous tumor-infiltrating T lymphocytes (TILs), which are expanded *ex vivo* and reinfused into patient, achieved success rates of over 30% and durable clinical benefits in patients with advanced melanoma refractory to ICB. These promising results for ACT-TIL led to accelerated FDA approval of lifileucel (Amtagvi®)^2,3^.

Unfortunately, many patients still do not respond, and there is further need for improvement of immunotherapies in solid tumors. The lack of response is most commonly due to insufficient persistence and exhaustion of the infused ACT-TIL product. CD8^+^ TILs, critical for anti-tumor immune responses overall, as well as for responses to ICB and ACT-TIL, have been shown to progress over time to a terminally exhausted state (Ttex), including in settings of ACT, due to chronic exposure to tumor antigens and a suppressive tumor microenvironment^4–7^. Ttex CD8^+^ TILs express high levels of multiple inhibitory receptor^i^s, including PD-1, TIM-3, TIGIT, LAG-3 and CTLA-4, proliferate poorly and cannot be reprogramed by ICB^8–10^. Several TFs have been shown to control the Ttex state, including Tox, Tox2, Nfat, Nr4a1 and Nr4a2^11–17^. Ttex CD8^+^ TILs differentiate from a progenitor population, termed Tpex, which express high levels of the stemness-associated TFs Tcf1, Lef1 and Bach2^18–21^ and the chromatin remodeling factor Satb1 ^22,23^, as well as several receptors, including Slamf6, Il7ra, Sell, Ccr7, Cxcr5 and CD28^8,24^. They have limited expression of inhibitory receptors, elevated proliferative capabilities and can be reprogrammed by ICB^4,25^. Hence, pathways that promote Tpex over Ttex states may potentiate CD8^+^ T cell anti-tumor responses, including in ACT TIL product.

The transcription factor Bcl11b controls T cell development at several stages, including at early commitment, as well as at beta and double positive selection stages^26–29^. In mature CD8^+^ T cells, during response to intracellular pathogens, conditional *Bcl11b* deletion, prior to activation, impaired TCR signaling and expansion^30,31^, while later, post-activation, conditional *Bcl11b* deletion potentiated differentiation of tissue resident memory (T_RM_) cells, including their cytolytic program^32^. A T_RM_ signature in the TILs is associated with positive outcomes, including in treatments with checkpoint inhibitors^33–35^.

Given the elevated T_RM_ program in the absence of *Bcl11b* in *Listeria* infection, in this study we investigated the role of Bcl11b in CD8^+^ T cells in anti-tumor response using both mouse models and human ACT-TILs. We show that deletion of *Bcl11b* in tumor Ag-specific CD8^+^ T cells results in major reduction in tumor burden in adoptive cell transfer models. *Bcl11b* KO CD8^+^ TILs show broad reprograming, having reduced exhaustion, but increased stemness. We found that Bcl11b controls expression of critical genes of these programs, repressing expression of TF genes associated with stemness, including Tcf7, Lef1, Bach2 and Satb1, and, conversely, promoting expression of exhaustion-associated TFs and inhibitory receptor genes. Additionally, our data show that Ag-specific *Bcl11b* KO CD8^+^ T cells kill more efficiently in an MHC class I-dependent manner and show increased Gzmb and Prf1 levels. CRISPR-mediated deletion of BCL11B in human TILs from a patient with poor response to ACT-TIL resulted in increased cytolytic activity, with elevated PRF1 and maintenance of high levels of the stemness-associated TF TCF1, suggesting that BCL11B may represent a therapeutic target to improve ACT-TIL.

## Results

### Low *BCL11B* mRNA is associated with improved survival of patients undergoing ICB for melanoma treatment

Recently we found that deletion of *Bcl11b* resulted in increased generation of intestinal T_RM_ CD8^+^ T cells following oral infection with *Listeria* in mice^32^. Presence of cells with a T_RM_ signature has been associated with improved prognosis in several cancers^36–39^. We thus analyzed data from the iATLAS Pan-ICI, which contains several studies of melanoma patients treated with ICB^40–44^, and the Moffitt melanoma study, also having patients treated with ICB^45^. *BCL11B* transcription normalized to T cell abundance, based on *CD3E* transcript levels was correlated with survival, using the PATH-SURVEYOR^46^. In both cohorts we observed that patients with improved overall survival exhibited lower *BCL11B* transcript levels relative to T cell abundance prior to treatment (**Fig. 1A-B**). We further analyzed a publicly available scRNA-seq dataset of basal cell carcinoma patients treated with ICB^47^. Our analysis revealed that CD8⁺ T cells of patients with complete response (CR) displayed diminished *BCL11B* mRNA levels compared to CD8⁺ T cells of non-responders, while mRNAs for the transcription factors associated with stemness BACH2 and SATB1 were elevated (**Fig. 1C**). These data suggest that diminished *BCL11B* expression in T cells, including in CD8^+^ TILs, may favor tumor control, providing a rationale for targeting *BCL11B* in T cell immunotherapy.

**Figure 1.**
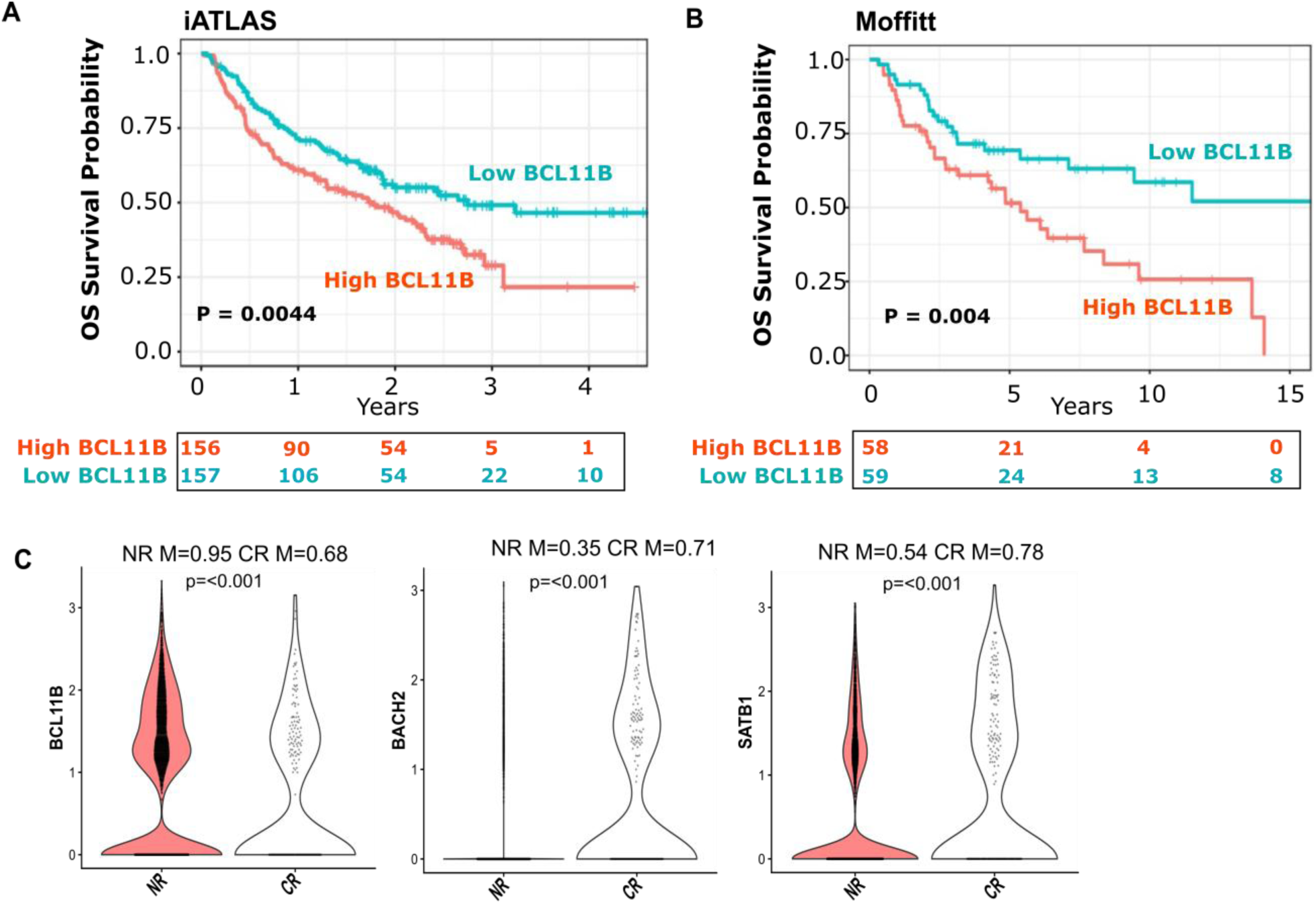
Patients with improved overall survival following check point inhibitor blockade (ICB) exhibit low *BCL11B* levels in T cells, including CD8^+^ T cells. **(A-B)** Association of *BCL11B* expression, normalized to T-cell abundance, and overall survival in melanoma patients treated with ICB. *BCL11B* and *CD3E* expression were converted to rank-based enrichment scores using the GSVA package (ssGSEA method). The difference between the two scores was calculated to estimate *BCL11B* expression relative to T-cell abundance using the PATH-SURVEYOR ^46^. Gene expression was evaluated prior to ICB therapy in (A) iATLAS (n = 313) and (B) Moffitt (n = 117) cohorts. iATLAS Hazard Ratio of 1.57 (95% CI: 1.15-2.14) and Moffitt Hazard Ratio of 2.20 (95% CI: 1.27-3.81). Kaplan–Meier survival curves of patients treated with ICB were categorized into high (red) and low (blue) *BCL11B*. **(C)** *BCL11B, BACH2* and *SATB1* mRNA levels in CD8^+^ T cells of patients with basal cell carcinoma with compete response (CR) versus non-responders (NR) to ICB treatment^47^. Gene expression is based on Log-normalized expression of UMI counts. CD8⁺ T cells were identified based on *CD3D/CD3E* and *CD8A/CD8B* expression. A statistical test was performed based on a Wilcoxon rank sum test. The mean expression for the CR and NR groups is indicated.

### Deletion of *Bcl11b* in CD8^+^ T cells post activation confers enhanced anti-tumor response in multiple solid tumor models

To investigate how BCL11B controls CD8^+^ T cell response in tumor immunity, we employed an adoptive cell transfer system in which splenic CD8^+^ T cells from PMEL *HGZMBCre*/*Bcl11b^f/f^*(KO) or PMEL *HGZMBCre* (WT) mice were pre-activated and expanded *ex vivo* with the PMEL specific gp100 peptide, and next transferred in mice bearing flank B16F10 tumors (**Fig. S1A**). *HGZMBCre* only becomes active post CD8^+^ T cell activation^48^, thus allowing to bypass the defects associated with TCR activation in the absence of *Bcl11b*^30,31^. YFP reporter showed efficient activity in the *HGZMBCre*/*Bcl11b^f/f^* system and removal of Bcl11b was efficient (**Fig. S2A-B**).

Transferred *Bcl11b* KO PMEL CD8^+^ T cells delayed B16F10 tumor growth compared to the transferred WT PMEL CD8^+^ T cells (**Fig. 2A**). Similarly, mice bearing flank B16F10-OVA tumors, transferred with ex vivo pre-activated and expanded *Bcl11b* KO OT-I CD8^+^ T cells (**Fig. S1B**), had smaller tumors, progressing slower compared to mice transferred with WT OT-I CD8^+^ T cells (**Fig. 2B**). Mice receiving Bcl11b-KO PMEL CD8^+^ T cells or Bcl11b-KO OT-I CD8^+^ T cells achieved a longer end-point threshold than those receiving WT OT-I CD8^+^ T cells or WT PMEL CD8^+^ T cells, respectively (**Fig. 2C, D**), suggesting that transfer of Bcl11b-KO CD8^+^ T cells may confer increased survival.

**Figure 2.**
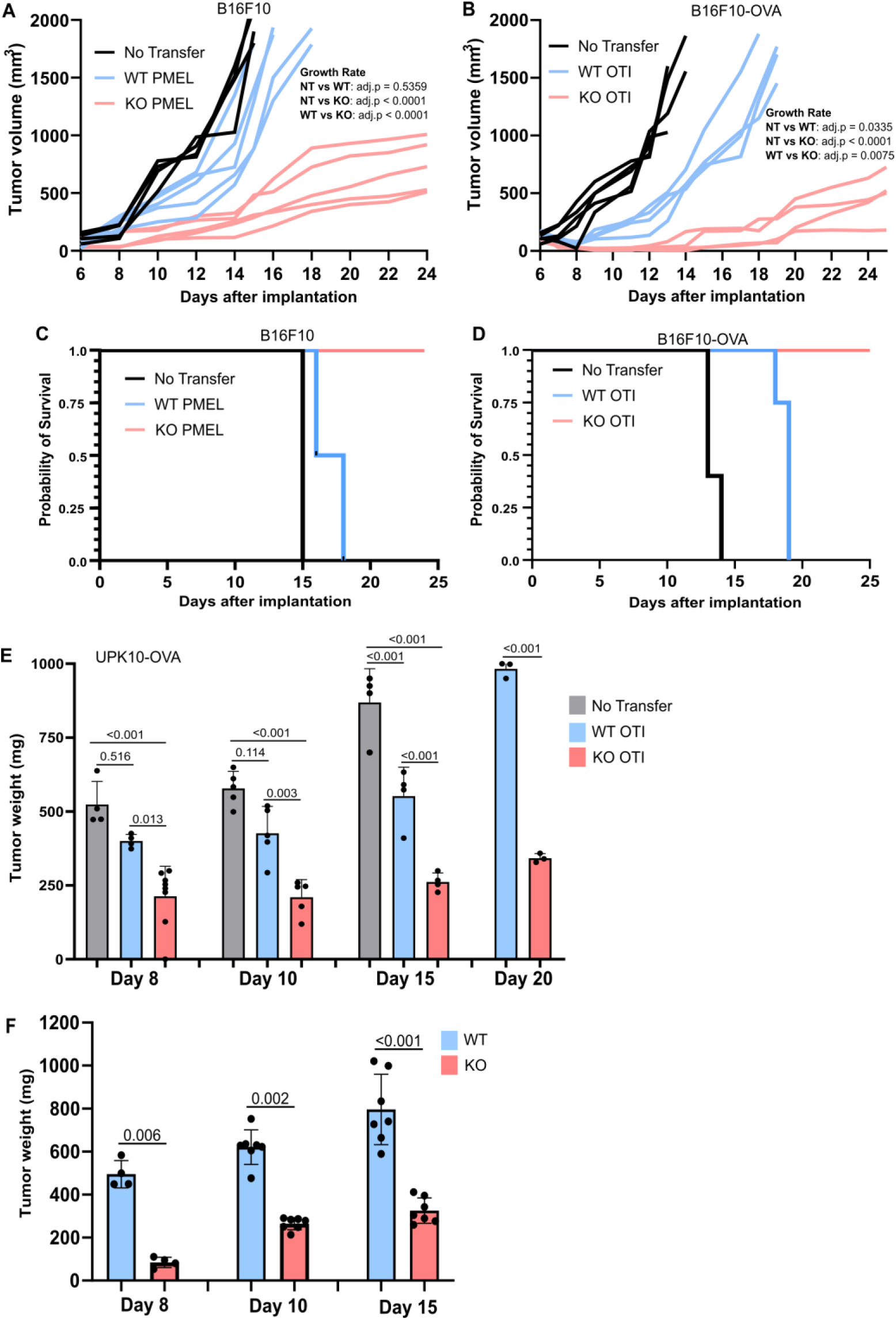
*Bcl11b* KO CD8^+^ TILs promote enhanced anti-tumor response in melanoma and ovarian tumor models. **(A)** Tumor volumes of CD45.1 mice injected subcutaneously (s.c.) on flank with melanoma B16F10 cells and transferred with PMEL CD8^+^ T cells derived from PMEL/*HGZMBCre*/*Bcl11b^f/f^*R26R-EYFP/CD45.2 (KO) or PMEL/ *HGZMBCre*R26R-EYFP/CD45.2 (WT) mice. **(B)** Tumor volumes of CD45.1 mice implanted on flank with B16F10-OVA cells and transferred with OT-I CD8^+^ T cells derived from OT-I/*HGZMBCre*/*Bcl11b^f/f^*R26R-EYFP/CD45.2 (KO) or OT-I/*HGZMBCre*R26R-EYFP/CD45.2 (WT) mice. (A-B) Statistical analysis for tumor growth (log2-transformed tumor size) was modeled using a linear mixed-effects model with fixed effects for time (Days), experimental group, and their interaction (growth slope), and a random intercept to account for subject variation. *p* values were derived from pairwise comparisons of tumor growth slopes performed using estimated marginal trends. (**C-D**) Kaplan-Meier survival analysis based on end point related to tumors of ≥2000 mm³ in PMEL B16F10 transfer system (C) and OT-I B16F10-OVA transfer system (D). **(E)** Weights of peritoneal tumors on Days 8, 10, 15 and 20. CD45.1 mice were intraperitoneally (i.p.) injected with UPK10-OVA ovarian tumor cells and transferred with OT-I CD8^+^ T cells derived from OT-I/*HGZMBCre*/*Bcl11b^f/f^*R26R-EYFP/CD45.2 (or CD45.1/2) (KO) or OT-I/*HGZMBCre*R26R-EYFP/CD45.2 (or CD45.1/2) (WT) mice. No Transfer are CD45.1 recipient mice with tumors but no transferred cells. **(F)** Weights of peritoneal tumors on Days 8, 10 and 15 in *HGZMBCre*/*Bcl11b^f/f^*R26R-EYFP (KO) and *HGZMBCre*R26R-EYFP/CD45.2 (WT) mice directly inoculated i.p. with UPK10 ovarian tumor cells. (E-F) Statistical analysis was performed using a two-way ANOVA followed by Tukey’s multiple-comparison test. Adjusted *p* values are shown. Experiments were conducted as detailed in **Fig. S1** and Material and Methods, including for activation of transgenic cells prior to transfer. “No transfer” represents CD45.1 mice inoculated with tumor cells, but not transferred with any transgenic CD8^+^ T cells. Data are representative of two independent experiments with n = 3-5 mice per group.

We employed an additional solid tumor model, namely the p53/Kras-driven ovarian carcinosarcoma UPK10 and UPK10-OVA, with cells injected intraperitoneally (i.p.)^39,49^ (**Fig. S1C, D**). Previously it was shown that in this model CD8^+^ T cells play an important role in anti-tumor immunity^39,50^. Results show that females with peritoneal UPK10-OVA tumors, transferred with *Bcl11b* KO OT-I CD8^+^ T cells, developed smaller tumors as early as Day 8 post-tumor inoculation, and further on Days 10, 15 and 20, compared to WT OT-I CD8^+^ T cells-transferred females, and with much slower progression (**Fig. 2E**).

In addition, *HGZMBCre*/*Bcl11b^f/f^* (KO) females directly injected i.p. with UPK10 cells, also developed significantly smaller tumors compared to WT females on Days 8, 10 and 15, and also with slower progression (**Fig. 2F**). These results together demonstrate that ablation of *Bcl11b* in CD8^+^ TILs post-activation reduces tumor burden more efficiently than WT.

### Analysis of heterogeneity and states of the transferred tumor *Bcl11b* KO and WT CD8^+^ TILs reveals increased divergence as the cells progress to the Ttex state

To establish the mechanisms that confer an advantage to *Bcl11b* KO CD8^+^ TILs in anti-tumor response, we conducted scRNAseq on transferred *Bcl11b* KO and WT OT-I CD8^+^ TILs in mice with peritoneal UPK10-OVA tumors. Unsupervised clustering analysis of transferred OT-I CD8^+^ TILs enabled the identification of 6 transcriptionally distinct clusters. The clusters were further defined based on specific and differential expression of genes known to be associated with stemness, including *Tcf7*, *Lef1*, *Bach2*, *Satb1*, *Runx1* and *Il7r*, versus exhaustion-associated genes, including *Nr4a1*, *Nr4a2*, *Pdcd1* and *Ctla-4* **(Fig. S3A)**. Principal component analysis (PCA) and Uniform Manifold Approximation and Projection (UMAP) were then utilized to evaluate TIL state heterogeneity in *Bcl11b* KO and WT CD8^+^ TILs samples (**Fig. 3A-D, S3B**). PC3 effectively separated *Bcl11b* KO versus WT CD8^+^ TILs (**Fig. 3A**). Two Tpex clusters (C0, C1) and one Ttex cluster (C5) were identified, as well as three additional intermediary clusters (Tinex) (C2, C3 and C4) with intermediate or low levels of expression of Tpex and Ttex genes **(Fig. S3A)**. Projection of the identified clusters onto the PCA space revealed that PC1 tracks the transition from stemness to exhaustion in both *Bcl11b* KO and WT CD8^+^ TILs (**Fig. 3B-C, S3B**). Further analysis revealed more similarities between *Bcl11b* KO and WT CD8^+^ TILs in the earlier Tpex states and more divergence in the later Ttex states, further confirmed by RNA velocity analysis (**Fig. 3B-D**). Notably, a gradual transition of more unspliced transcripts from Tpex clusters toward more spliced transcripts in the Tinex3 and Ttex clusters was observed (**Fig. S3B**), in line with the more differentiated state of latter clusters.

**Figure 3.**
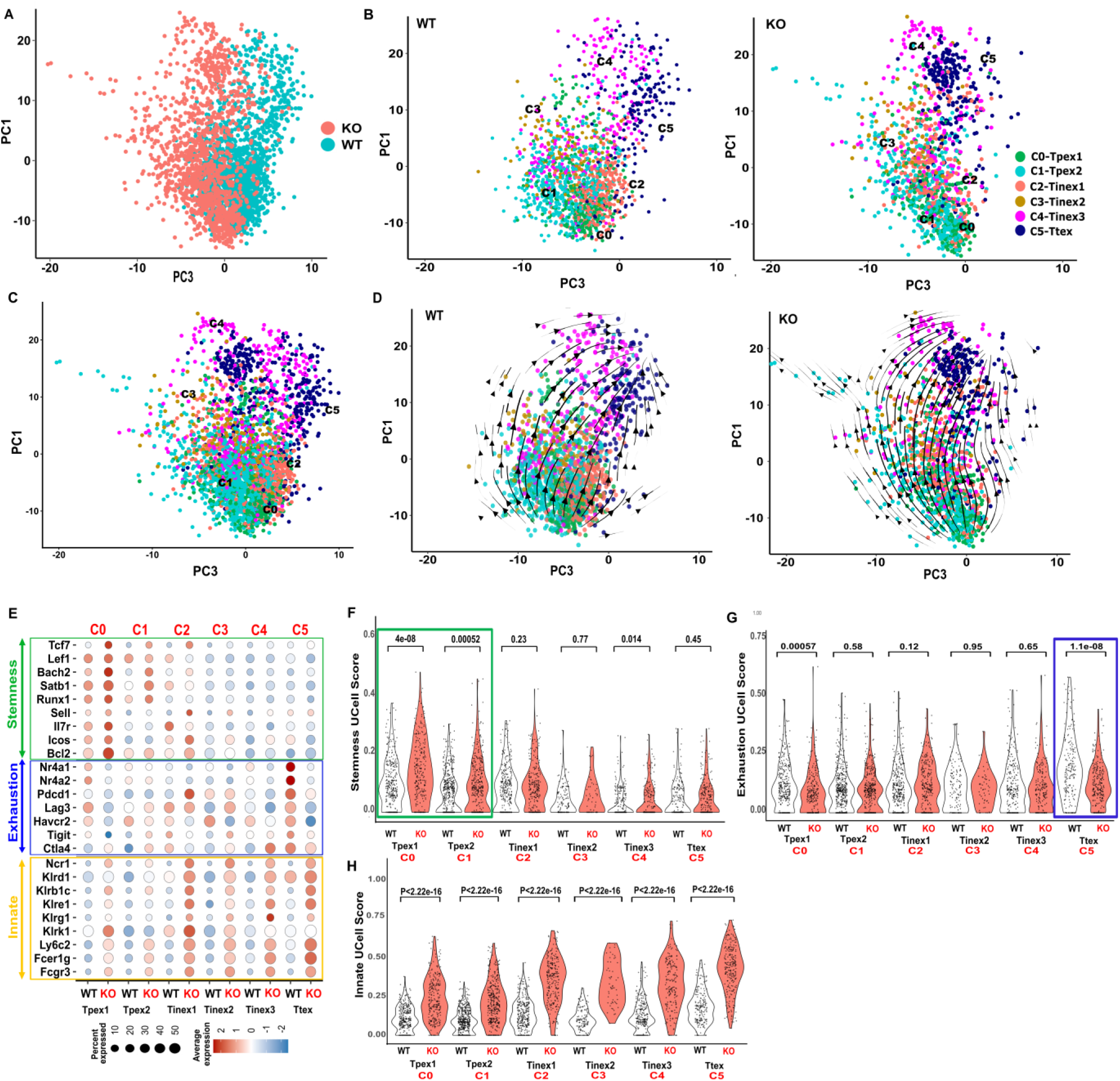
*Bcl11b* KO CD8⁺ TILs upregulate stemness program and show limited progression to the Ttex state. scRNA-seq analysis of transferred OT-I CD8⁺ T cells isolated from peritoneal UPK10-OVA tumors. CD45.1 mice were injected i.p. with UPK10-OVA ovarian tumor cells and transferred with CD8^+^ T cells from donor CD45.2 OT-I *HGZMBCre*/*Bcl11b^f/f^*R26R-EYFP or *HGZMBCre*R26R-EYFP mice, activated as described in Fig. S1C. **(A)** Principal component analysis (PCA) of scRNA-seq of transferred OT-I CD8⁺ T cells isolated from peritoneal UPK10-OVA tumors. PC1 is associated with differentiation states and PC3 with the analyzed group (*Bcl11b* KO and WT). **(B)** Clustering analysis of transcriptionally distinct T cell states, annotated as: progenitors of exhausted T cells (Tpex1, C0, and Tpex2, C1), intermediary exhausted (Tinex1, C2, Tinex2, C3, Tinex3, C4), and terminally exhausted (Ttex, C5). **(C)** PCA projection of integrated clustering of transferred *Bcl11b* KO and WT OT-I CD8^+^ TILs. **(D)** Velocity analysis using UniTVelo reveals directional lineage trajectories, supporting progressive differentiation from Tpex toward Ttex states. (**E**) Dot heatmap showing differential expression of genes associated with stemness, exhaustion and innate programs in transferred *Bcl11b* KO and WT OT-I CD8^+^ TIL clusters. **(F-H)** Violin plots derived from UCell algorithm analysis showing scores for stemness **(F)**, exhaustion **(G)** and innate **(H)** programs. Statistical significance was assessed using the Wilcoxon rank-sum test.

Thus, *Bcl11b* KO and WT CD8^+^ TILs are heterogeneous, progress from Tpex to Ttex states and show more pronounced divergence in the Ttex state.

### Stemness program is elevated, while exhaustion program is reduced in *Bcl11b* KO CD8^+^ TILs

We further investigated differential gene expression between transferred *Bcl11b* KO and WT CD8^+^ TILs, focusing on stemness and exhaustion programs. Tpex1 (C0) cluster showed the highest increase in stemness signature gene expression in the absence of *Bcl11b*, followed by Tpex2 (C1), exemplified by elevated expression of *Tcf7*, *Lef1*, *Bach2*, *Satb1*, *Runx1*, *Sell*, *Il7r*, *Icos and Bcl2* (**Fig. 3E**). Supporting this observation, Ucell algorithm stemness score was significantly higher in the *Bcl11b* KO Tpex1 and Tpex2 clusters compared to the WT (**Fig. 3F**). Thus, in the context of anti-tumor immunity, opposite from the previous observations in the *Bcl11b* KO T_RM_ CD8^+^ T cells in the response to oral infection with Listeria^32^, *Bcl11b* KO CD8^+^ TILs upregulate the stemness program.

The WT Ttex cluster C5 showed the highest levels of genes associated with exhaustion, including for the TFs Nr4a1 and Nr4a2, together with several genes encoding inhibitory receptors, including Pd-1, Lag-3, Tim-3, Tigit and Ctla-4, all strikingly low in the *Bcl11b* KO C5 (**Fig. 3E**), also supported by the Ucell score (**Fig. 3G**). These results suggest a failure of the *Bcl11b* KO TILs to progress to the Ttex state.

Similar to previous findings in *Bcl11b* KO T_RM_ CD8^+^ T cells, as well as in the CD4^+^ T cells and Treg cells, but not in NK cells^32,51–53^, innate genes, including *Fcer1g* and *Fcgr3*, as well as NK receptor genes, including several Klr family members, were upregulated in *Bcl11b* KO CD8^+^ TILs in all clusters (**Fig. 3E, H**).

Altogether, these results reveal the extensive transcriptional reprogramming induced by *Bcl11b* deletion in CD8^+^ TILs. Stemness-associated genes are elevated in the absence of *Bcl11b*, and there is a pronounced reduction of exhaustion signature genes in the Ttex cluster, overall suggesting that *Bcl11b* KO CD8^+^ TILs fail to progress to the Ttex state.

### *Bcl11b* KO CD8^+^ TILs remain predominantly in the Tpex state

Given the reprogramming of TILs following *Bcl11b* deletion, we further conducted flow cytometry analysis using combinations of different markers for evaluation of Tpex and Ttex subsets in the transferred OT-I CD8^+^ T cells in UPK10-OVA tumors.

The Tpex subset, defined as Slamf6^hi^PD-1^lo^, CD62^hi^Lag3^lo^, CD127^hi^Tim-3^lo^ or Tcf1^hi^Tox^lo^ was highly enriched in the transferred *Bcl11b* KO CD8^+^ TILs, while the Ttex subset, defined as Slamf6^lo^PD-1^hi^, CD6L^lo^Lag3^hi^, CD127^lo^Tim-3^hi^ or Tcf1^lo^Tox^hi^, was reduced, opposite from the distribution in the transferred WT CD8^+^ TILs (**Fig. 4A-D**). In addition, a Bach2+ population was predominant in the transferred *Bcl11b* KO CD8^+^ TILs and poorly represented in the WT (**Fig. 4E**). Confirming scRNAseq data, these results demonstrate that transferred *Bcl11b* KO CD8^+^ TILs remain predominantly in a Tpex state, with limited progression to the Ttex state.

**Figure 4.**
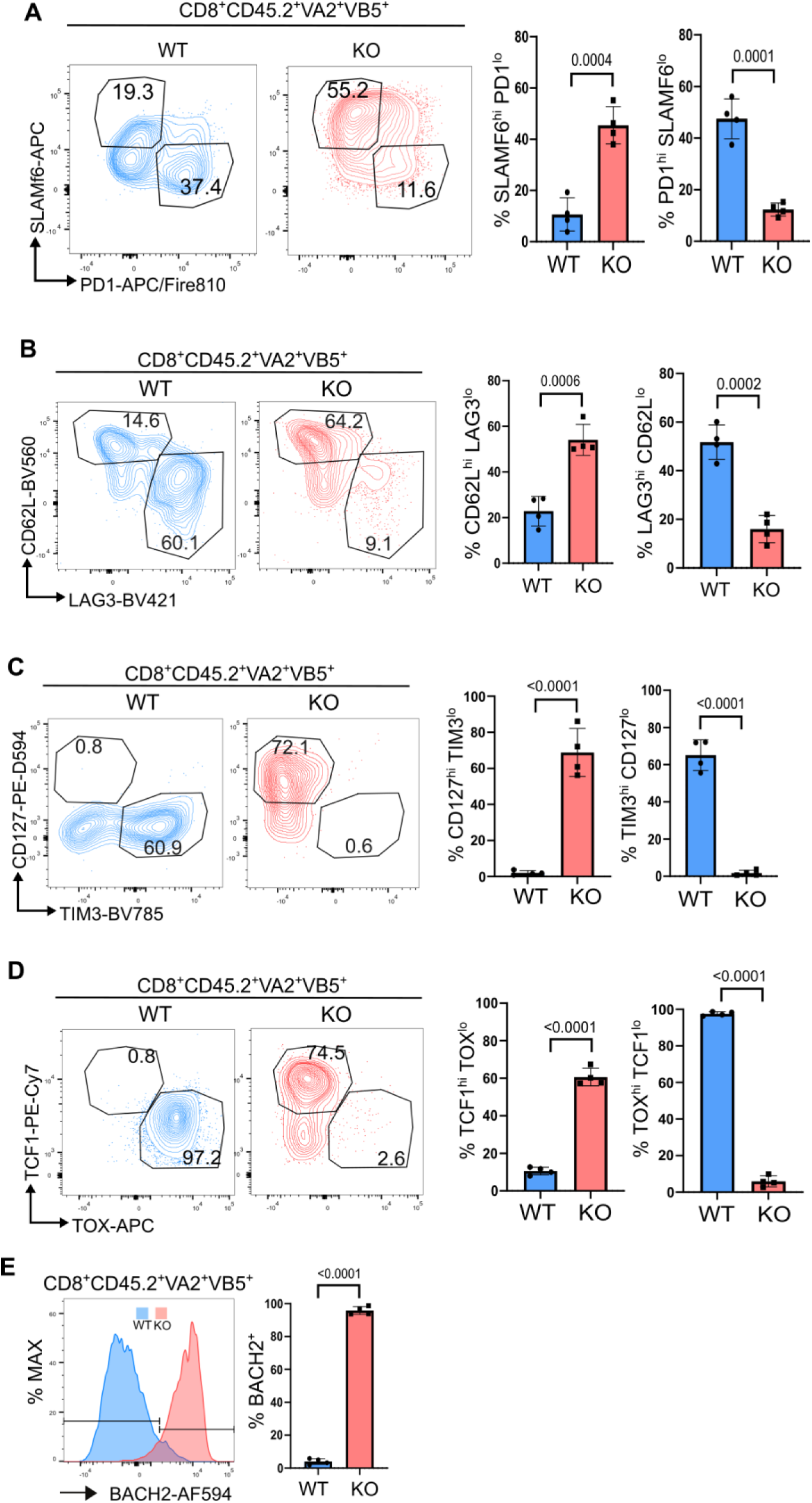
Tpex CD8^+^ TILs predominate following *Bcl11b* deletion. Adoptive transfers of CD8^+^ T cells derived from donor CD45.2 OT-I *HGZMBCre*/*Bcl11b^f/f^*R26R-EYFP (KO) or *HGZMBCre*R26R-EYFP (WT) mice into CD45.1 mice transplanted intraperitoneally (i.p.) with UPK10-OVA as described in **Fig. S1C**. Before transfer CD8^+^ T cells were activated as described in **Fig. S1** and in Material and Methods. **(A-D)** Representative contour plots and statistics of the stemness vs exhaustion subsets, defined by the indicated markers, in the gated transferred TCRβ+CD8+CD45.2+Vα2+Vβ5+ *Bcl11b* KO and WT CD8^+^ TILs on Day 10 after UPK10-OVA tumor inoculation. **(A)** Slamf6 vs PD1, **(B)** CD62L vs Lag3, **(C)** CD127 vs TIM-3, **(D)** TCF1 vs TOX. **(E)** Representative histogram for Bach2 in TCRβ+CD8+CD45.2+Vα2+Vβ5+ CD8^+^ TILs. Data are representative for two independent experiments with n = 3-4 mice per group. *p* values by unpaired t test.

### *Bcl11b* KO CD8^+^ T cells promote increased antigen-specific killing restricted to MHC class I+ targets

Given the reduction in tumor burden in the absence of *Bcl11b* in CD8^+^ TILs and the central role of cytotoxicity in tumor control, we next evaluated the functionality of *Bcl11b* KO and WT CD8^+^ T cells in terms of killing in an Ag-dependent manner, specifically of UPK10-OVA and B16F10-OVA tumor cells. *Bcl11b* KO OT-I CD8^+^ T cells showed elevated killing activity against both UPK10-OVA and B16F10-OVA tumor cells compared to WT (**Fig. 5A-F**). However neither *Bcl11b* KO nor WT CD8^+^ T cells (OT-I or polyclonal) killed the MHC class I-negative YAC-1 cells (**Fig. 5A-C**, **Fig. 5G-I**), which are known targets for NK cells^54^, despite the fact that *Bcl11b* KO CD8^+^ TILs upregulated several NK receptor mRNA genes, as well as surface NK1.1 and NKp46 (**Fig. 3E**, **Fig. 5J**). These results demonstrate that *Bcl11b* KO CD8^+^ T cells have elevated killing activity and only kill in an antigen MHC class I-dependent manner.

**Figure 5.**
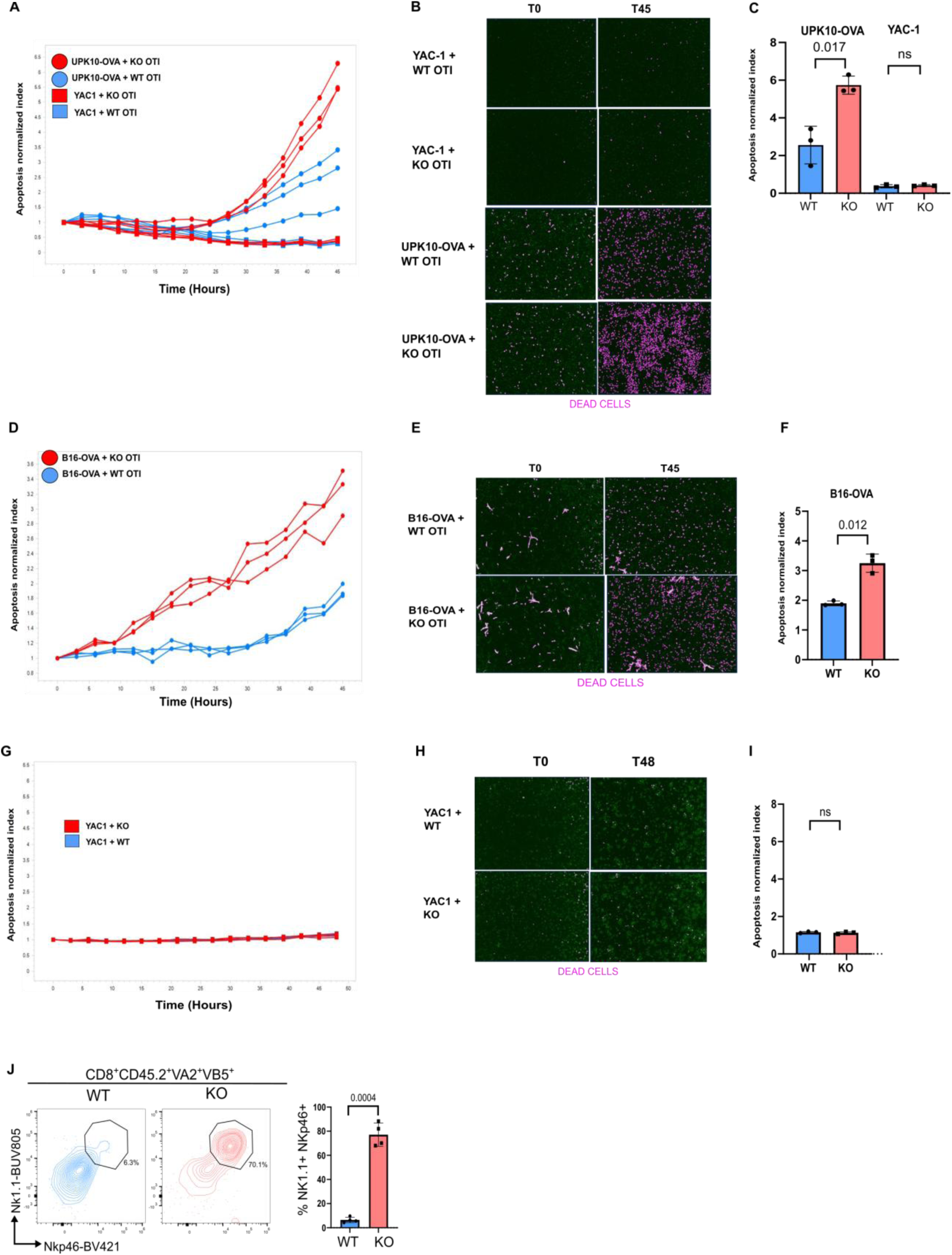
*Bcl11b* KO CD8^+^ TILs have increased Ag-specific MHC class I-dependent cytolytic activity. **(A-F)** Splenocytes from OT-I/*HGZMBCre*/*Bcl11b^f/f^*R26R-EYFP or OT-I/*HGZMBCre*R26R-EYFP mice were activated ex vivo and purified, as indicated in **Fig. S1** and Material and Methods. **(A-C)** The activated CD8^+^ OT-I cells were co-cultured with UPK10-OVA or YAC-1 or **(D-F)** with B16F10-OVA, for 45 hrs in 96-well plates in the presence of Caspase 3/7 Green Dye. Details are provided in Material and Methods. Images were recorded on IncuCyte SX5 (Sartorius AG) every 3 hours for 45 hours. **(B, E)** Images at 45 hrs. An overlay mask (magenta color) is shown on the images to indicate cells counted as positive for the apoptotic reagent. **(C, F)** Apoptosis normalized index at 45 hrs. **(G-I)** Preactivated polyclonal CD8^+^ T cells from *HGZMBCre*/*Bcl11b^f/f^*R26R-EYFP or *HGZMBCre*R26R-EYFP mice were co-cultured with YAC-1 cells as in (A), in the presence of Caspase 3/7 Green Dye. Tumor cell apoptosis was recorded every 3 hours for 48 hours and data was analyzed as in (A-F). **(I)** Apoptosis normalized index at 48 hrs. **(J)** Flow cytometry analysis of Nk1.1 and Nkp46 in transferred *Bcl11b* KO and WT OT-I CD8^+^ TILs (TCRb+CD8+CD45.2+Vα2+Vβ5+) isolated from peritoneal UPK10-OVA tumors of recipient CD45.1 mice. Adoptive transfers were performed as described in Fig. S1C. Analysis was conducted on day 10 after tumor inoculation. *p* values by unpaired t test.

### *Bcl11b* KO CD8^+^ TILs have elevated Gzmb and Prf1 and maintain high TCF1

Given the elevated cytolytic activity of *Bcl11b* KO CD8^+^ T cells compared to the WT counterparts, we evaluated Gzmb and Prf1 proteins in transferred Ag-specific *Bcl11b* KO and WT CD8^+^ TILs in the UPK10-OVA tumors. The results show increased Gzmb and Prf1 proteins in OT-I *Bcl11b* KO CD8^+^ TILs, compared to the WT counterparts (**Fig. 6A**). However neither *Gzmb* nor *Prf1* mRNAs were upregulated in the *Bcl11b* KO CD8^+^ TILs (**Fig. 6B-C**). Prf1 and Gzmb proteins have been previously shown to be post-transcriptionally regulated and their levels could increase during response without changes in transcription^55^. In line with this, the cytoplasmic translation pathway was elevated in *Bcl11b* KO C4 (Tinex3) and C5 (Ttex) TIL clusters (**Fig. S4A-D**). Expression of numerous ribosomal genes was elevated in *Bcl11b* KO C4 (Tinex3) and C5 (Ttex) TIL clusters, together with that of genes encoding the eIF3 translation initiation factor protein complex, including *Eif3i*, *Eif3j1* and *Eif3m* (**Fig. S4C-D**). Additional genes upregulated only in the C5 cluster included those encoding several subunits of eukaryotic translation elongation factor 1 (eEF1) complex, namely *Eef1a1*, *Eef1b2* and *Eef1g* (**Fig. S4D**). Thus the increase in Prf1 and Gzmb proteins in *Bcl11b* KO CD8^+^ TILs, without elevation in their mRNAs, may be associated with elevated translational activity.

**Figure 6.**
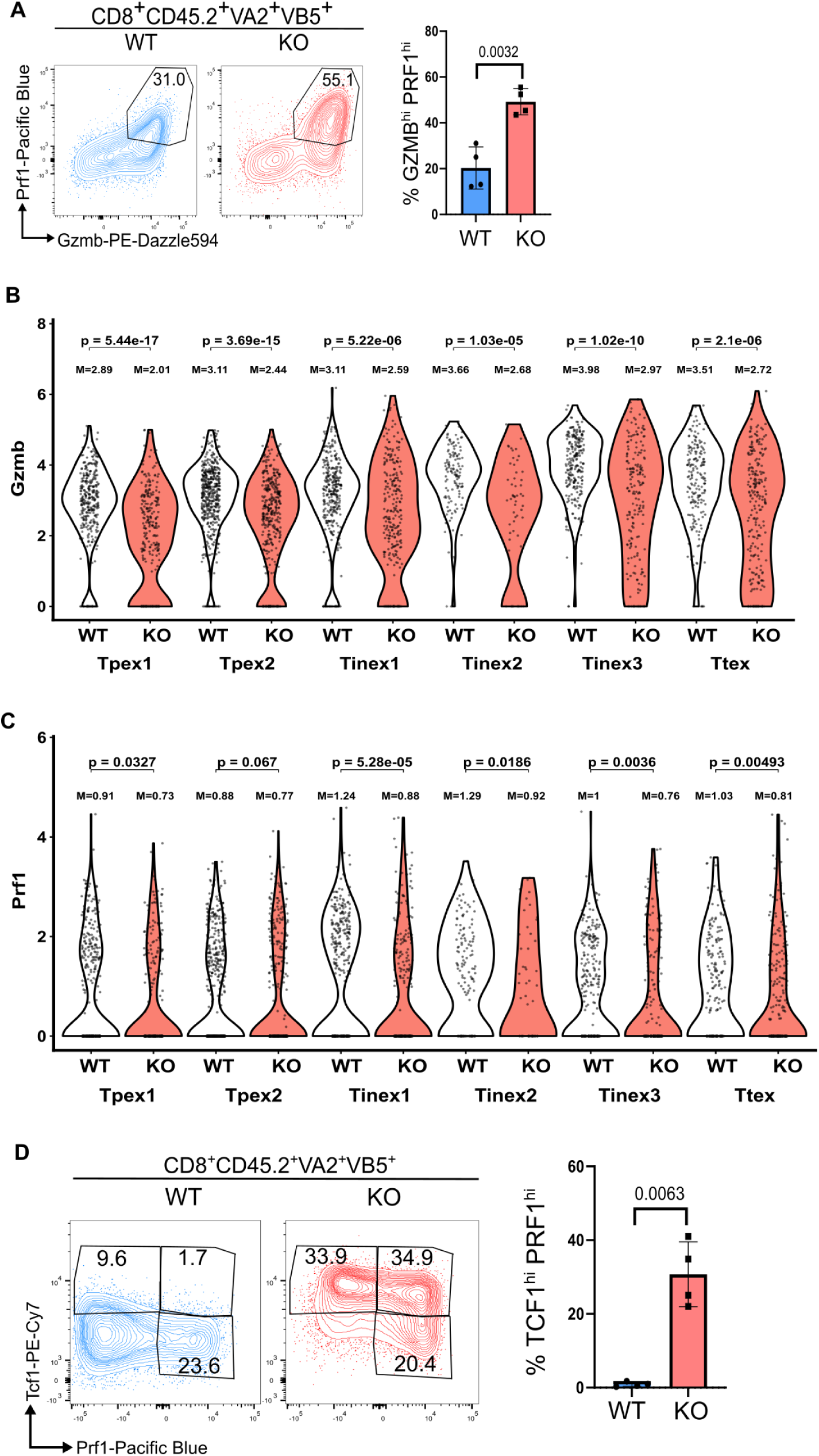
*Bcl11b* KO CD8^+^ TILs show elevated Prf1 and Gzmb together with high TCF1. **(A)** Flow cytometry analysis of Prf1 and Gzmb in transferred *Bcl11b* KO and WT OT-I CD8^+^ TILs (TCRb+CD8+CD45.2+Vα2+Vβ5+) isolated from peritoneal UPK10-OVA tumors of CD45.1 mice. Adoptive transfers were performed as described in Fig. S1C. Analysis was conducted on day 10 after tumor inoculation. *p*-value by unpaired *t*-test. Data are representative of two independent experiments. **(B, C)** Violin plots showing Gzmb and Prf1 mRNA across clusters, stratified by genotype. *p* value are based on the Wilcoxon rank-sum test assessing the differential expression between groups. Mean expression values for each group are indicated on the plots based on log normalized mRNA. **(D)** Flow cytometry analysis of Prf1 and Tcf1 in transferred *Bcl11b* KO and WT OT-I CD8^+^ TILs (TCRb+CD8+CD45.2+Vα2+Vβ5+) isolated from peritoneal UPK10-OVA tumors of CD45.1 mice as in (A). *p*-value by unpaired *t*-test.

We show that *Bcl11b* KO CD8^+^ TILs have increased Tcf1 (**Fig. 4D**). Tcf1 was previously shown to repress expression of *Prf1*^56^, which is in agreement with the reduced *Prf1* mRNA levels in *Bcl11b* KO CD8^+^ TILs (**Fig. 6C**). However a fraction of *Bcl11b* KO CD8^+^ TILs high in Tcf1, still show elevated Prf1 protein (**Fig. 6D**), further suggesting a post-transcriptional regulatory mechanism related to elevated Prf1. Thus *Bcl11b* KO CD8^+^ TILs maintain both elevated Tcf1 and Prf1, possessing multipotency and high cytolytic activity.

### Bcl11b-dependent regulation of the epigenetic landscape of CD8^+^ TILs

To uncover the mechanisms by which Bcl11b controls transcriptional programs and epigenetic landscape in TILs, we performed **C**leavage **U**nder **T**argets and **R**elease **U**sing **N**uclease (CUT&RUN) ^57^ and **A**ssay for **T**ransposase-**A**ccessible **C**hromatin (ATAC)-seq to characterize Bcl11b binding in the genome, along with the deposition of the histone marks H3K27ac and H3K27me3, and chromatin accessibility (ChrAcc), respectively, on polyclonal CD8^+^ TILs isolated from peritoneal UPK10-OVA tumors. H3K27ac is a prominent histone mark associated with transcriptional activation, including with enhancer activity, while H3K27Me3 is a well-established histone mark for gene silencing^58^. Deletion of *Bcl11b* increased H3K27Ac and chromatin accessibility signals both at promoters and enhancers bound by Bcl11b (**Fig. 7A-D**), suggesting that Bcl11b acts predominantly through repression of target genes, likely dependent on the NuRD complex, as we previously published^59^, NuRD complex is known to mediate repression through histone deacetylase activity (HDAC1/HDAC2)^60^. We further ranked H3K27Ac stitching using the ROSE algorithm to establish putative super-enhancers (SEs)^61,62^. In both WT and *Bcl11b* KO CD8^+^ TILs, SEs were found at several genes associated with exhaustion, including *Tox*, *Nr4a2, Pdcd1* and *Ctla-4* (**Fig. 7E**). However, only in the *Bcl11b* KO CD8^+^ TILs, putative SE activity was also present at the stemness associated genes *Tcf7, Lef1, Bach2* and *Satb1* (**Fig. 7E**), in agreement with their elevated expression in *Bcl11b* KO CD8^+^ TILs. *Tcf7* and *Bach2* genes showed broad enrichment of H3K27Ac, together with elevated chromatin accessibility in *Bcl11b* KO CD8^+^ TILs, while *Satb1* and *Lef1* genes showed both increased H3K27Ac, as well as a broad reduction in H3K27Me3 (**Fig. 7F-I**). Additional genes with elevated H3K27Ac signals in *Bcl11b* KO TILs included ribosomal and translation genes (*Rpl11, Rpl28, Rpl36* and *Eif3i)* (**Fig. S5A-D**).

**Figure 7.**
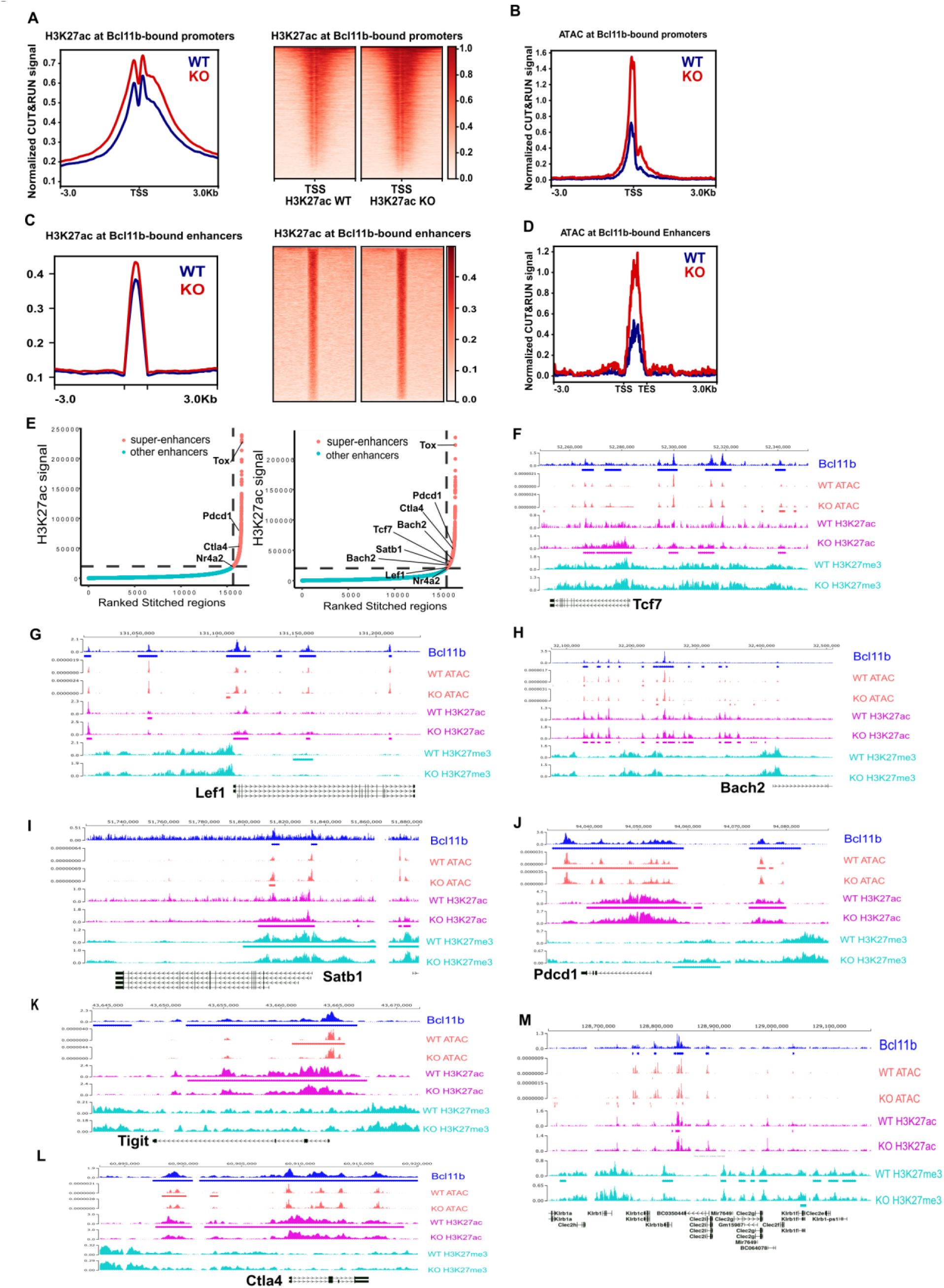
Epigenetic landscape controlled by Bcl11b in CD8^+^ TILs. (**A**) H3K27ac signal at transcription start site (TSS ±3 kb) at Bcl11b-bound promoters. **(B)** ATAC-seq signal at TSS (±3 kb) at Bcl11b-bound promoters. **(C)** H3K27ac signal at Bcl11b-bound enhancers. **(D)** ATAC-seq signal at Bcl11b-bound enhancers. **(E)** Super-enhancers versus typical enhancers defined by H3K27Ac signal intensity and stitching over 12.5kb by Rose Algorithm. **(F-M)** Genomics tracks for ATAC-seq and CUT&RUN for Bcl11b, H3K27Ac and H3K27me3 at the indicated genes. Rectangles beneath each track indicate peaks that are significantly different between WT and *Bcl11b* KO cells, or significantly bound by Bcl11b. CUT&RUN-seq for H3K27ac, H3K27me3, as well as ATACseq were conducted on polyclonal CD8^+^ TILs isolated from peritoneal UPK10-OVA tumors of *HGZMBCre*/*Bcl11b^f/f^*R26R-EYFP (KO) and *HGZMBCre*R26R-EYFP/CD45.2 (WT) mice. CUT&RUN-seq for Bcl11b was conducted on polyclonal CD8^+^ TILs isolated from peritoneal UPK10-OVA tumors of WT mice. Details are provided in Material and Methods, including for the analysis.

Conversely, at several genes encoding inhibitory receptors, including *Pdcd1, Ctla-4* and *Tigit*, there was a decrease in H3K27ac signal, together with reduction in chromatin accessibility in *Bcl11b* KO TILs (**Fig. 7J-L**). This points to more complex and discrete regulation of target gene expression mediated by Bcl11b, through the use p300 histone acetyl transferase, to promote transcriptional activation^63^, despite the overall increase in H3K27Ac and ChrAcc at promoters and enhancers bound by Bcl11b (**Fig. 7A-D**). Noticeably, *Pdcd1* also showed increased H3K27Me3 (**Fig. 7J**). Related to H3K27Me3, a remarkable and broad reduction of this mark occurred at the NK gene complex on chromosome 6 in the absence of *Bcl11b* (**Fig. 7M**), which also correlated with increased ChrAcc, but no changes in H3K27Ac. The broad reduction in H3K27Me3 and increased ChrAcc were associated with upregulation of numerous *Klr* gene expression, located in this genomic region (**Fig. 3E, 3I, 7M**). At other derepressed innate genes, such as *Fcer1g*, there was a remarkable increase in H3K27Ac associated with ChrAcc, but no difference in H3K27Me3 (**Fig. S5E**), suggesting that Bcl11b may employ diverse mechanisms to repress expression of innate genes.

These results collectively demonstrate that deletion of *Bcl11b* results in epigenetic reprogramming at several key genes with roles in stemness, exhaustion and innate genes.

### Deletion of *BCL11B* in human TILs from a patient with poor response to ACT-TIL conferred increased cytolytic activity and elevated TCF1

Given the positive impact of *Bcl11b* deletion in CD8^+^ TILs in adoptive cell transfers in both ovarian and melanoma murine models, we investigated the impact of *BCL11B* deletion by CRISPR/Cas9-editing in human CD8^+^ TILs from a melanoma patient with poor response to ACT-TIL therapy. gBCL11B-CRISPR-edited TILs showed reduction of BCL11B versus control, but no change in CD8 or CD3, while the NK receptor CD56 was upregulated (**Fig. 8A-D**), in line with the derepression of the NK receptor genes observed in the mouse TILs. In addition, *BCL11B* KO TILs had elevated PRF1 and TCF1 continued to express PRF1 in TCF1^hi^ cells (**Fig. 8E**), similar to what was observed in the mouse CD8^+^ TILs. We further tested the cytolytic activity of the TILs against HLA-matched tumor cells. *BCL11B* KO TILs displayed elevated cytolytic activity against tumors compared to the gCTRL-CRISPR-edited TILs (**Fig. 8F-G**). These results showed that *BCL11B* deletion in TILs promotes elevated anti-tumor cytolytic activity, likely associated with increased levels of PRF1. Thus, BCL11B may constitute a potential therapeutic target to improve T cell fitness in immunotherapeutic settings of solid tumors.

**Figure 8.**
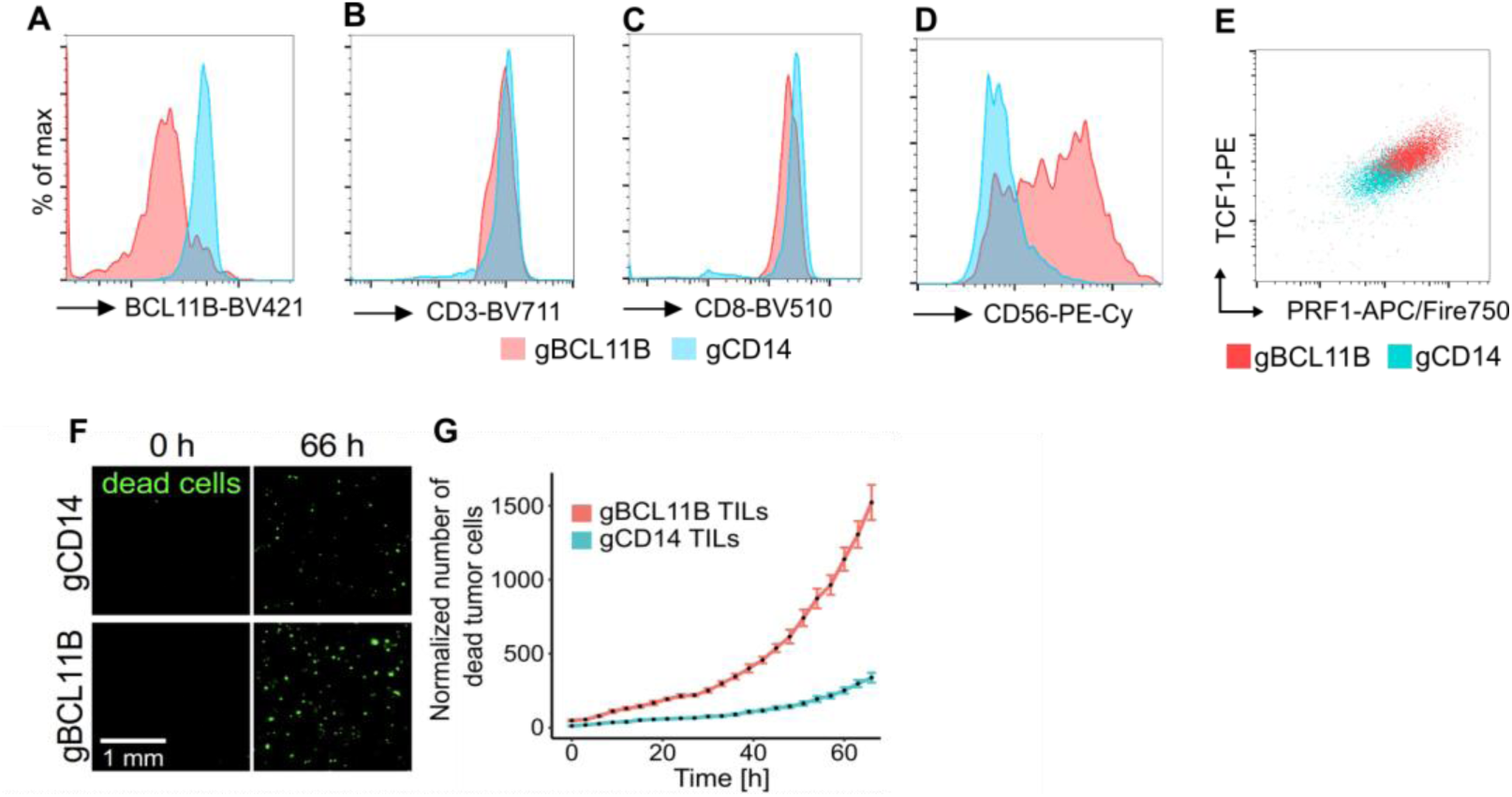
Depletion of BCL11B from human melanoma TILs from a patient with poor response to ACT-TIL leads to elevated cytotoxicity and increased TCF1. **(A-E)** TILs derived from a non-responder patient to ACT-TIL were CRISPR/Cas9-edited with gBCL11B or gCD14 (control). CD14 is not expressed in human CD8^+^ T cells. Flow cytometry analysis of BCL11B (A), CD3 (B), CD8 (C), CD56 (D) and TCF1 vs PRF1 (E) in TILs CRISPR/Cas9-edited with gBCL11B or gCD14 (control). (**F-G**) Cytolytic activity of *BCL11B* KO versus control TILs. The killing activity was evaluated against patient-derived MHC-I matched tumor cells. Dead cells were stained with Caspase 3/7 Green detection reagent added to the medium as described in Material and Methods.

## Discussion

In this study we found that in the absence of *Bcl11b* CD8^+^ TILs promoted a reduced tumor burden in solid tumor murine models. Furthermore, *BCL11B* deletion by CRISPR-editing in human ACT-TILs conferred elevated cytolytic response with increased PRF1 and the stemness TF TCF1, findings most impressive as the TILs were derived from a patient who responded poorly to ACT-TIL. In the murine model, *Bcl11b* KO CD8^+^ TILs were broadly reprogrammed, predominantly remaining in the Tpex and intermediate states with poor progression to the Ttex state. Our results demonstrate that Bcl11b is positioned upstream of critical genes of the stemness and exhaustion programs, blocking expression of the former and sustaining expression of the latter. Thus Bcl11b deletion results in concomitant upregulation of stemness program and reduction of exhaustion program, offering an advantage in anti-tumor response of CD8^+^ TILs. *Bcl11b* KO Tpex TILs upregulated stemness associated TF genes, including *Tcf7, Lef1* and *Bach2*, as well as the chromatin organizer and transcriptional regulator Satb1 gene, recently found to be important for stem-like CD8^+^ progenitors and Tpex CD8^+^ T cells in chronic infection, and to control expression of *Tcf7, Bach2* and *Myb*^22,23^. The increased stemness program in *Bcl11b* KO CD8^+^ TILs is opposite to our previous findings in CD8^+^ T_RM_ cells in response to oral infection with Listeria, where we found that deletion of *Bcl11b* resulted in reduction of *Tcf7* and overall of the stemness program^32^. This points to a context dependent activity of *Bcl11b* in CD8^+^ T cells in tumors versus acute infection. Importantly in both models we depleted *Bcl11b^f/f^* with the same HGZMBCre deleter. Our results show that Bcl11b binds at *Tcf7, Lef1, Bach2* and *Satb1* genes. At *Tcf7* and *Bach2* genes, absence of *Bcl11b* resulted in increased H3K27Ac activity and chromatin accessibility. At *Satb1* and *Lef1* genes, both increase in H3K27Ac was observed, together with reduction in H3K27Me3. Bcl11b bound at several inhibitory receptor genes, including *Pdcd1, Tigit* and *Ctla4*, and absence of Bcl11b predominantly reduced H3K27Ac. Similarly to these findings, Bcl11b was previously found to control H3K27ac to regulate expression of specific target genes in Treg cells, MAIT cells and memory CD8^+^ T cells ^32,52,64^. H3K27Ac was found increased in the case of upregulated genes, or reduced for downregulated genes, which is in line with the previous finding that BCL11B associates with the NuRD complex, involved in transcriptional repression^59^, or with the p300 histone acetyl transferase, known to promote active gene expression^63^.

Interestingly, Satb1 was previously shown to use the NuRD complex to silence *Pdcd1*^65^. It is possible that at least in part, the impact on *Pdcd1* is related to the increased Satb1 expression. However, Bcl11b also bound at the *Pdcd1* locus, suggesting a direct control. *Bcl11b* deletion not only caused elevated H3K27Ac at *Pdcd1* gene, but also increased H3K27Me3, suggesting additional mechanisms of silencing.

A broad reduction of H3K27Me3 was observed at the NK gene complex on chromosome 6 in the absence of *Bcl11b*, correlated with increased ChrAcc, suggesting major reprograming of the region, associated with broad upregulation of expression of numerous *Klr* genes. Nonetheless, *Bcl11b* KO CD8^+^ T cells did not kill YAC-1 cells, known targets for NK cells, and only killed in an MHC class I-dependent manner. Upon a more detailed analysis of the upregulated NK receptor genes, we noticed that not only activating receptors, *Klrb1c, Klrk1* – located in the NK gene complex on chromosome 6, and *Ncr1*, located on chromosome 7, were increased in expression, but inhibitory receptor genes as well, including *Klrg1, Klre1* and *Klrd1*. This may explain the failure of *Bcl11b* KO CD8^+^ T cells to kill YAC-1 cells, while maintaining the ability to kill in an antigen-specific manner, which is critically important for ACT with autologous TILs. Even if *Bcl11b* KO CD8^+^ T cells did fail to kill the MHC class I negative cells YAC-1, it is still possible that derepression of the activating NK receptors may potentiate the killing activity of *Bcl11b* KO CD8^+^ T cells.

Interestingly, deletion of *Bcl11b* did not result in increased H3K27Ac at the NK complex genes, but predominantly broad reduction of the H3K27Me3 mark, suggesting that Bcl11b may function at this genomic region through mechanisms independent of the NuRD complex, and potentially involving EZH2 of the PRC2 complex, shown recently to repress expression of NK receptor genes in human CAR-T cells by deposition of H3K27Me3^66^.

At other derepressed genes, such as *Fcer1g*, no changes in H3K27Me3 were observed, however H3K27Ac was elevated, suggesting a possible dependence on the NuRD complex for transcriptional repression.

Related to increased Gzmb and Prf1 proteins in *Bcl11b* KO CD8^+^ TILs, but not elevated mRNAs, we found that expression of numerous ribosomal genes was elevated together with that of genes encoding the translation initiation factor protein complex eIF3, as well as of the translation elongation complex eEF1. Thus the increase in Prf1 and Gzmb may be related to the elevated translational activity, given that they have been previously shown to accumulate without changes in transcription ^55^.

The changes in the programs in the absence of *Bcl11b* in CD8^+^ TILs were reflected also in subset distribution as evaluated by flow cytometry analysis. Remarkably, a fraction of BCL11B KO TILs with elevated PRF1 levels continued to express TCF1 both in murine and human TILs, despite that *TCF7* and effector genes typically display dissociated expression^67^. These results suggest that *BCL11B* KO CD8^+^ TILs retain multipotency, and that targeting of BCL11B in TILs boosts cytotoxicity and stemness, promoting an enhanced anti-tumor response.

In conclusion, our investigations suggest that downregulation of *BCL11B* in T cells is not only correlated with improved survival of melanoma patients treated with ICB, but deletion of *BCL11B* in TILs from a patient with poor response to ACT-TIL improved TIL activity and likely retained multipotency. In mouse models, we demonstrate that *Bcl11b* deletion in TILs alters cell states, promoting increased cytotoxic capacity, yet blunting exhaustion and paradoxically increasing stemness. These findings establish an important role for BCL11B in regulating mature CD8^+^ T cell responses to solid tumors and highlight *BCL11B* as a potential target for improvement of T cell therapies against solid tumors.

## RESOURCE AVAILABILITY

### Lead contact

Information and requests should be directed to the lead contact Dorina Avram. All materials and resources will be made available upon request.

### Data availability

Single-cell RNA-seq, ATAC-seq and CUT&RUN-seq data associated with this study have been deposited in the Gene Expression Omnibus (GEO) database with accession number GSE327609, GSE327438 and GSE327416.

Code: https://github.com/shawlab-moffitt/AvramLab_BCL11B_CD8Tcells_CodeRepo

## ACKNOWLEDGMENTS

Studies were supported by R01CA293755 (to D.A. and T.I.S.), R01AI067846 (to D.A.), Adelson Medical Research Foundation (G-202206-00524) (to P.H.), Swim Across America and Ocala Royal Dames for Cancer Research (to S.P-T.) and P30-CA076292 (to Moffitt Cancer Center). We thank Dr. Dung-Tsa Chen for help with statistical data analysis for murine tumor progression.

## AUTHOR CONTRIBUTIONS

Conceptualization: L.S., T.Z., V.B.C., D.P.T., S.P-T. and D.A; data curation: L.S., D.P.T, T.Z, V.B.C., A.N.O., T.I.S. and D.A.; formal analysis: L.S., D.P.T, T.Z, V.B.C., S.I., R.P.S.,T.I.S. and D.A.; investigation: L.S., T.Z, D.P.T, V.B.C., S.I., R.P.S., Z.N., S.C.; methodology: L.S., T.Z., D.P.T, S.I., R.P.S., V.B.C., J.R.C-G., S.Z.M, W.H., Y.T.B., ML.S.H., J.L.B, S.P-T., resources: J.R.C-G., S.Z.M., P.H., E.E., ML.S.H., J.L.B, S.P-T., J.O.J., A.S., A.A.T., J.E.M., E.D.; visualization: L.S., D.P.T, T.Z., V.B.C., J.O.J. and D.A.; supervision: D.A., T.I.S., S.P-T., funding acquisition: D.A.; project administration; DA; writing – original draft: L.S., T.Z., V.B.C., D.P.T, T.I.S., S.P-T. and D.A.

## Materials and Methods

### Mice

*Bcl11b^f/f^* mice, previously described ^27,29,30,51,52,64,68–71^, were crossed to B6-Tg(*GZMB*-cre)1Jcb/J (*HGZMBCre*) mice, that have the Cre transgene cloned under the control of the human *GZMB* promoter^48^, and to the R26R-EYFP Cre reporter mice^72^. For adoptive transfers models, mice were further crossed to PMEL^73^ or OT-I^74^ mice, all from Jackson Laboratory. All mice are on C57BL/6. B6.SJL-PtprcaPepcb/BoyJ (CD45.1) recipient mice were purchased from Charles River Laboratory, bred in our colony and used as recipients for adoptive transfers. Mice of both sexes were used at age between 7–12 weeks, except for UPK10 ovarian tumors, where females were used both as recipients and donors. All animals were maintained by the animal facility of the Moffitt Cancer Center, housed in a temperature controlled (18-23°C), 40-60% humidity, 12 h light/dark cycle facility. Animal studies were performed in accordance with the Institutional Animal Care and Use Committee of the University of South Florida Research Integrity and Compliance department. Animal Welfare Assurance Number A4100-01.

### Cell lines

UPK10 cells, which are p53/Kras-driven ovarian carcinosarcoma^49^ were transduced to express Ovalbumin (OVA) and sorted for low OVA expression^39^. B16F10-OVA was generated similarly.

### Mouse CD8^+^ T-cells activation and treatment for adoptive transfers

Splenocytes from naïve OT-I or PMEL mice were grown in MEM alpha with 10% FBS, 1% penicillin-streptomycin, 1% sodium pyruvate, 1% nonessential amino acids and 50 μM β-mercaptoethanol, plus 100 U/ml IL2 and 1 μg/mL SIINFEKEL OT-I peptide or 1 μg/mL PMEL peptide (MMOTOPES) for 48 hours at 37°C, 5% CO2, when cells were split 1:3 in fresh medium with IL-2, for an additional 72 hours, when they were transferred in mice with tumors or used in killing assays. For polyclonal CD8^+^ T cells splenocytes were activated in the same conditions, plus 1 μg/mL anti-CD3 and 2 μg/mL anti-CD28 antibodies (BioXCell) and further used in killing assays.

## Tumor Models

### Adoptive transfer - Melanoma flank models

Naïve CD45.1 mice were injected s.c. on flank with 0.5×10^6^ B16F10 or B16F10-OVA melanoma cells. Mice were further transferred i.v. with 1×10^6^ CD8^+^ T cells from donor CD45.2 or CD45.1/2 PMEL or OT-I *HGZMBCre*/*Bcl11b^f/f^*R26R-EYFP or *HGZMBCre*R26R-EYFP mice. Prior to transfer, splenocytes from PMEL or OT-I mice were cultured in MEMa plus 10% FBS, 1% L-glutamine, 1% Pen/Strep, 50 μM β-mercaptoethanol and 100U IL2, plus PMEL KVPRNQDWL or OT-I SIINFEKL peptides, for 2 days, when peptide was removed, and cells were further cultured for 3 more days with IL2 only. Prior to transfer in mice with tumors on Day 7 post-tumor inoculation, CD8^+^ T cells were purified with anti-CD8 biotinylated antibodies and Mojort streptavidin nanobeads (Biolegend). Tumors were measured with an electronic caliper starting with Day 6, every other day.

### Adoptive cell transfer - Peritoneal tumor model

Naïve CD45.1 mice were injected i.p. with 10×10^6^ UPK10-OVA ovarian tumor cells and transferred on Day 5 post-tumor inoculation with 1×10^6^ CD8^+^ T cells from donor CD45.2 or CD45.1/2 OT-I *HGZMBCre*/*Bcl11b^f/f^*R26R-EYFP or *HGZMBCre*R26R-EYFP mice, activated as described above. Mice were euthanized on Days 8, 10, 15 and 20, when tumors were weighed. FACS analysis and scRNAseq were conducted on transferred TILs on Day 10 post-tumor inoculation.

### Peritoneal tumors in polyclonal mice

*HGZMBCre*/*Bcl11b^f/f^*R26R-EYFP and *HGZMBCre*R26R-EYFP mice directly injected i.p. with 10×10^6^ UPK10-OVA tumor cells. ATACseq and CUT&RUN for Bcl11b and histone marks were conducted on CD8^+^ TILs on Day 10 post-tumor inoculation.

### Culture and CRISPR of human TILs

The human melanoma TILs were previously expanded ex vivo in the presence of tumor cells and IL-2 and further selected for IFNG production and then then cryopreserved by the Pilon-Thomas lab ^75^. Cryopreserved TILs were thawed and cultured in complete media supplemented with 3000 U/ml IL-2 ^75^.

For CRISPR editing, after 2 days in culture, TILs were electroporated (3×10^5^/reaction) using Invitrogen™ Neon™ Transfection System. The Hs.Cas9.BCL11B.1.AA and Hs.Cas9.BCL11B.1.AE gRNAs^32^ (designed by IDT software) were combined, while Hs.Cas9.CD14.1.AA served as a negative control. Guide RNAs together with Alt-R® Cas9 Electroporation Enhancer, Alt-R® CRISPR-Cas9 tracrRNA, ATTO™ 550 and Alt-R S.p and Cas9 nuclease V3 (IDT) were assembled into RNP complexes and electroporated in cells in the following conditions: 1600 V, 10 ms, and 3 pulses. TILs were further cultured for 6 additional days before analysis.

### Murine TIL isolation and flow cytometry analysis

All tumor single-cell suspensions were obtained by tumor dissociation with the gentleMACS Dissociator (Miltenyi) and digestion with collagenase I (1 mg/mL), and DNAse I (100 ug/mL) for 15 min at 37 °C (only for B16F10). After red blood cell lysis, cells resuspended in FACS buffer (PBS buffer containing 2% FBS and 2 mM EDTA), further stained with fluorescence-labeled antibodies for 30 min at 4 °C. For staining of intracellular proteins, cells were washed in PBS and resuspended in fixation buffer for 30 min at 4 °, and then in permeabilization buffer and primary antibodies for 30 min at 4 °C. Samples were acquired on Aurora Cytek Cytometer and analyzed with FlowJo 10.10.1. Single stained cells and Fluorescence Minus One (FMO) sample were used as control. Antibodies and reagents used are listed in Supplementary Table 1.

### Human samples – TILs

The studies with human TILs were approved by the University of South Florida (USF) or Advarra Institutional Review Boards (IRB) under approvals Ame5_107905 (USF), Ame13_Pro00009061 (USF), and 14.03.0083 (Advarra), respectively, and informed written consent was received prior to participation.

### Cytotoxicity Assays. Mouse cells

For cytotoxicity assays, activated CD8^+^ T cells were cocultured with target cells: UPK10-OVA, B16F10-OVA or YAC-1 in 96-well plates, and 10000 tumor cells were incubated with 5000 CD8^+^ T cells in 100 μl of cell culture medium. Tumor cell apoptosis was detected by Caspase-3/7 Detection Reagent (Invitrogen) added to the medium. Data was acquired every 3 hours for 45 hours with Incucyte SX5 and analyzed by the IncuCyte software.

### Cytotoxicity Assays. Human cells

After six days in culture following CRISPR editing, 30,000 TILs were incubated with 5000 matched target tumor cells per well in 96-well plate. Tumor cell apoptosis was detected by Caspase-3/7 Detection Reagent added to the medium. Data was acquired every 3 hours for the duration of 69 hours with Incucyte SX5 and analyzed by the IncuCyte software.

### ATACseq library preparation and processing

ATAC-seq was performed as previously described ^32,76^ on 10,000 CD8^+^ TILs from peritoneal UPK10-OVA tumors. Briefly, nuclei were prepared by resuspending cells in 50 μl cold ATAC-RSB-NTD buffer and incubating for 3 minutes and then washed and incubated in 50 μl Transposition Mix, at 37°C for 30 minutes. Reactions were purified by Zymo DNA Clean and Concentrator-5 Kit and used for library preparation with the NEBNext High-Fidelity 2× PCR Master Mix and IDT for Illumina DNA/RNA UD indexes (Illumina). DNA was further purified using SPRIselect magnetic beads (Beckman Coulter) and sequenced as 2×150bp on NovaSeq 6000 or NovaSeq X platforms.

ATAC-seq was analyzed using PEPATAC ^76^ (v2.0.0) with default settings to generate fixed-size peaks with reduced biases by MACS2 ^76^ and Genrich ^77^. Differential analysis was performed by DESeq2 ^78^.

### CUT&RUN library preparation and processing

CUT&RUN was performed on CD8^+^ TILs from peritoneal UPK10-OVA tumors, 30,000 for H3K27me3; 100,000 for H3K27ac and 400,000 for Bcl11b. Samples were processed by the ChIC/CUT&RUN Kit (EpiCypher) with 1 µg of anti-H3K27me3 (Active Motif), 1 µg of anti-H3K27ac (EpiCypher), and combined 0.5 µg (#A300-385A, Bethyl) + 0.5 µg (#12120, D6F1, Cell Signaling Technology) anti-Bcl11b antibodies. For histone marks, we followed the manufacturer’s instructions. For Bcl11b, during the incubation step with protein A/G fused to micrococcal nuclease (pAG-MNase), cells were treated with eBioscience Perm Buffer (Invitrogen) supplemented with Spermidine and cOmpleteEDTA-free protease inhibitor (Roche). Libraries were prepared with the NEBNext Ultra II Library Prep Kit (New England Biolabs) and sequenced as 2×150bp on the Illumina NovaSeq 6000 or NovaSeq X platforms. Cells from one to two animals were pooled per sample to obtain enough cells.

For data processing, raw CUT&RUN reads were trimmed Trimmomatic (v0.39) and aligned to the mm10 or hg38 genome using Bowtie2 ^79^ (v2.3.5.1). Duplicates were removed using Picard (v2.25.5). Peaks were called by MACS2 (v2.2.5). Resulting bedgraph files were converted to the bigwig format by the bedGraphToBigWig tool from ENCODE-DCC/kentUtils and used for visualization. In parallel, peaks were called using SEACR ^80^ (v1.3) by selecting the top 1% of regions by AUC and using the stringent mode. Peaks from all biological replicates were concatenated, and overlapping peaks were merged across all the tested conditions by bedtools merge (v2.30.0)^81^. The resulting unique list of peaks was used for the differential analysis: raw counts for each peak were determined by bedtools coverage (v2.30.0), and these were analyzed using DESeq2 ^78^. The peaks were called differential when they had a *p* value < 0.05. Permissive differential peaks were also included with a combination of *p* value < 0.3 and |log2FC| > 0.2. Super enhancers were defined based on H3K27ac peaks ROSE (v1.3.2), stitched within 12.5 kb ^61^. Genome-wide analyses of binding patterns were performed using deepTools ^82^.

### Single-cell RNAseq library preparation and data processing

scRNAseq libraries were prepared by the PIPseqT2 3’ Single Cell RNA Kit v4.0PLUS (FBS-SCR-T2-8-V4.05, Fluent BioSciences) and PIPseq T2 3’ Single Cell RNA Kit v4.0 (FBS-SCR-T2-8-V4, Fluent BioSciences), according to the manufacturer’s instructions. Libraries were sequenced as 2×150bp on the Illumina NovaSeq 6000 or NovaSeq X platforms.

All samples were pre-processed by pipseeker (v3.3.0, Fluent Biosciences) with the reference files provided by GRCm39 (GENCODE vM29 2022.04, Ensembl 106). The processed feature, barcode, and matrix files were used as an input for our BERLIN pipeline^83^ for further processing and filtering, relying on Seurat (v5.0.1)^84^. Low-abundance genes/transcripts, doublet cells, and low-quality cells were removed prior to analysis. Data were transformed by SCTransform (v2)^85^ and used for integration of all replicates and conditions using FindIntegrationAnchors and IntegrateData function. Dimension reduction and clustering analysis were performed on integrated data. Identification of major T cell subset clusters was according to the expression of genes associated with stemness (Tcf7, Lef1, Bach2, Satb1, Sell, Runx1, Il7r, and Bcl2) and exhaustion (Nr4a1, Nr4a2, Pdcd1, LAG-3, Havcr2, Tigit, CTLA-4). Heterogeneous clusters were further subclustered at a higher resolution of 0.7, which was then used for final annotation. Clusters with fewer than ten cells or those that did not express signature T cell genes were removed.

The principal component analysis (PCA) was performed based on the unintegrated RNAseq data. To identify which PCs were most strongly associated with genotype, we compared PC scores between KO and WT samples for each PC using Welch’s two-sample t-test. The resulting t-statistics quantified the magnitude and direction of separation between genotypes along each PC. PCs were ranked by the absolute value of the t-statistics. PC3, with the largest t-statistics, was interpreted as capturing the dominant genotype-associated sources of variation.

The Wilcoxon rank sum test was used to identify differentially expressed genes in each cluster compared to those in all other clusters using the Seurat FindAllMarkers function. Similarly, the FindMarkers function was used to determine differential genes between two conditions. Gene set enrichment analysis (GSEA) was performed on pre-ranked differentially expressed genes using –log(*P* value) * sign(logFC) as ranks, using genetic signatures from the gene ontology (GO), the Molecular Signatures Database (MSigDB), the Kyoto Encyclopedia of Genes and Genomes (KEGG). Signature scores were calculated by the AddModuleScore function of UCell^86^, using the stemness and exhaustion signature gene sets.

For RNA velocity analysis, bam files from the pipseeker pipeline were converted to loom files using velocyto^87^, further combined with the metadata from the integrated Seurat object, and processed by UniTVelo (v0.2.5.2)^88^.

### Statistics

Statistical analyses were performed with Prism software (GraphPad), and an unpaired two tail T-test was used. *p* values of <0.05 were considered statistically significant. Peritoneal tumor weights were compared across genotype/treatment groups on Days 8, 10, 15, and 20 using a two-way ANOVA followed by Tukey’s multiple-comparison test. For flank tumor growth, a linear mixed-effect model was used with fixed effects for time (Days), experimental group, and their interaction (growth slope), and a random intercept to account for subject variation. *p* values were derived from pairwise comparisons of tumor growth slopes performed using estimated marginal trends.

---

**Figure S1.**
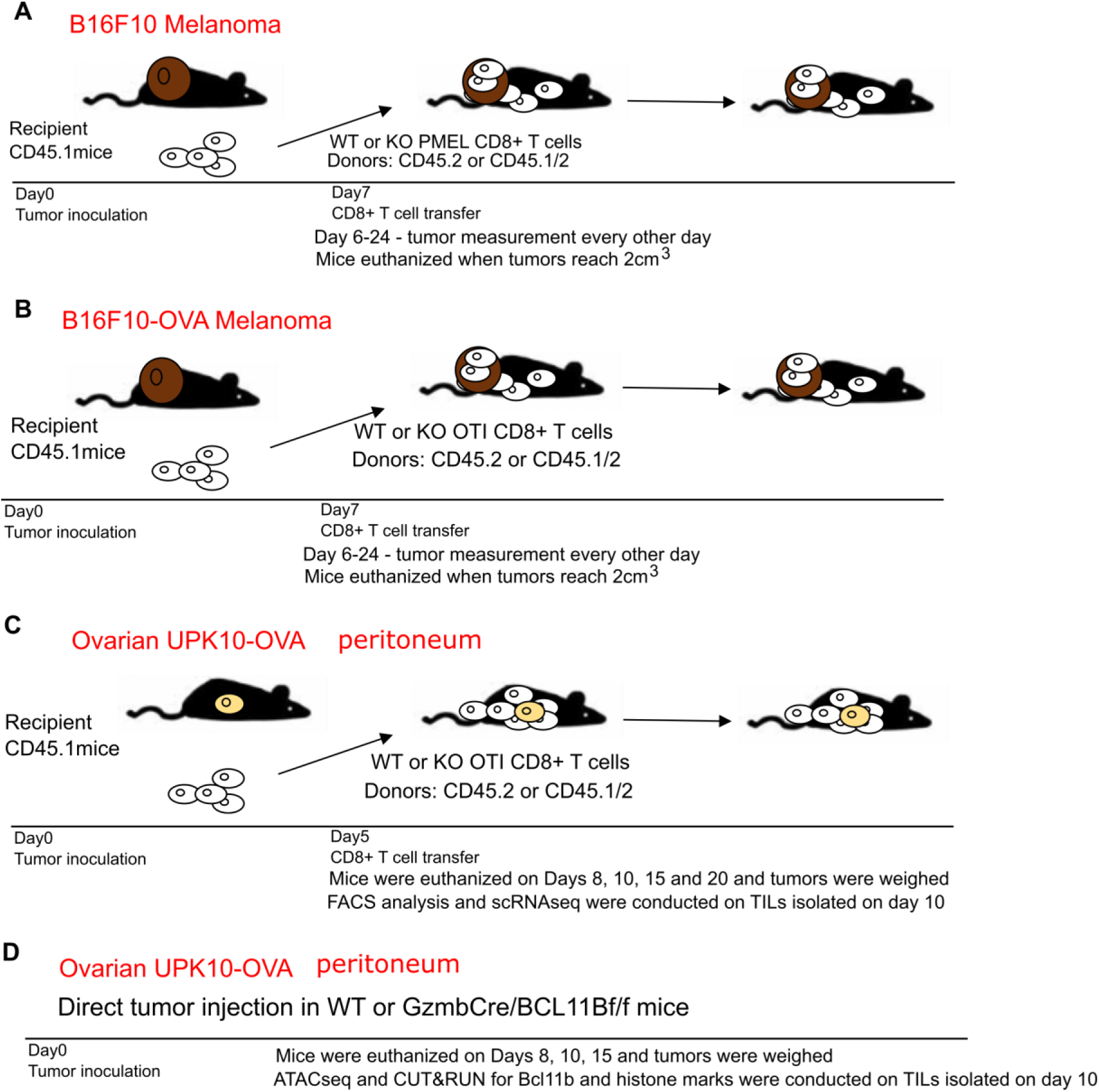
Experimental details for adoptive T cell transfers in murine models with tumors. **(A-B)** Adoptive cell transfer - melanoma flank models. Naïve CD45.1 mice were injected s.c. on flank with 0.5×10^6^ B16F10 (A) or B16F10-OVA (B) melanoma cells. Mice were further transferred i.v. with 1×10^6^ CD8^+^ T cells from donor CD45.2 or CD45.1/2 PMEL or OT-I *HGZMBCre*/*Bcl11b^f/f^*R26R-EYFP or *HGZMBCre*R26R-EYFP mice. Prior to transfer, splenocytes from PMEL or OT-I mice were cultured in MEMa plus 10% FBS, 1% L-glutamine, 1% Pen/Strep, 50 μM β-mercaptoethanol and 100U IL2, plus PMEL KVPRNQDWL or OT-I SIINFEKL peptides, for 2 days, when peptide was removed, and cells were further cultured for 3 more days with IL2 only. Prior to transfer, CD8^+^ T cells were purified with anti-CD8 biotinylated antibodies and Mojort streptavidin nanobeads (Biolegend). Tumors were measured with an electronic caliper. **(C)** Adoptive cell transfer - peritoneal model. Naïve CD45.1 mice were injected i.p. with 10×10^6^ UPK10-OVA ovarian tumor cells and transferred with 1×10^6^ CD8^+^ T cells from donor CD45.2 or CD45.1/2 OT-I *HGZMBCre*/*Bcl11b^f/f^*R26R-EYFP or *HGZMBCre*R26R-EYFP mice, activated as described in (A-B). **(D)** Polyclonal *HGZMBCre*/*Bcl11b^f/f^*R26R-EYFP and *HGZMBCre*R26R-EYFP mice directly injected i.p. with 10×10^6^ UPK10 tumor cells.

**Figure S2.**
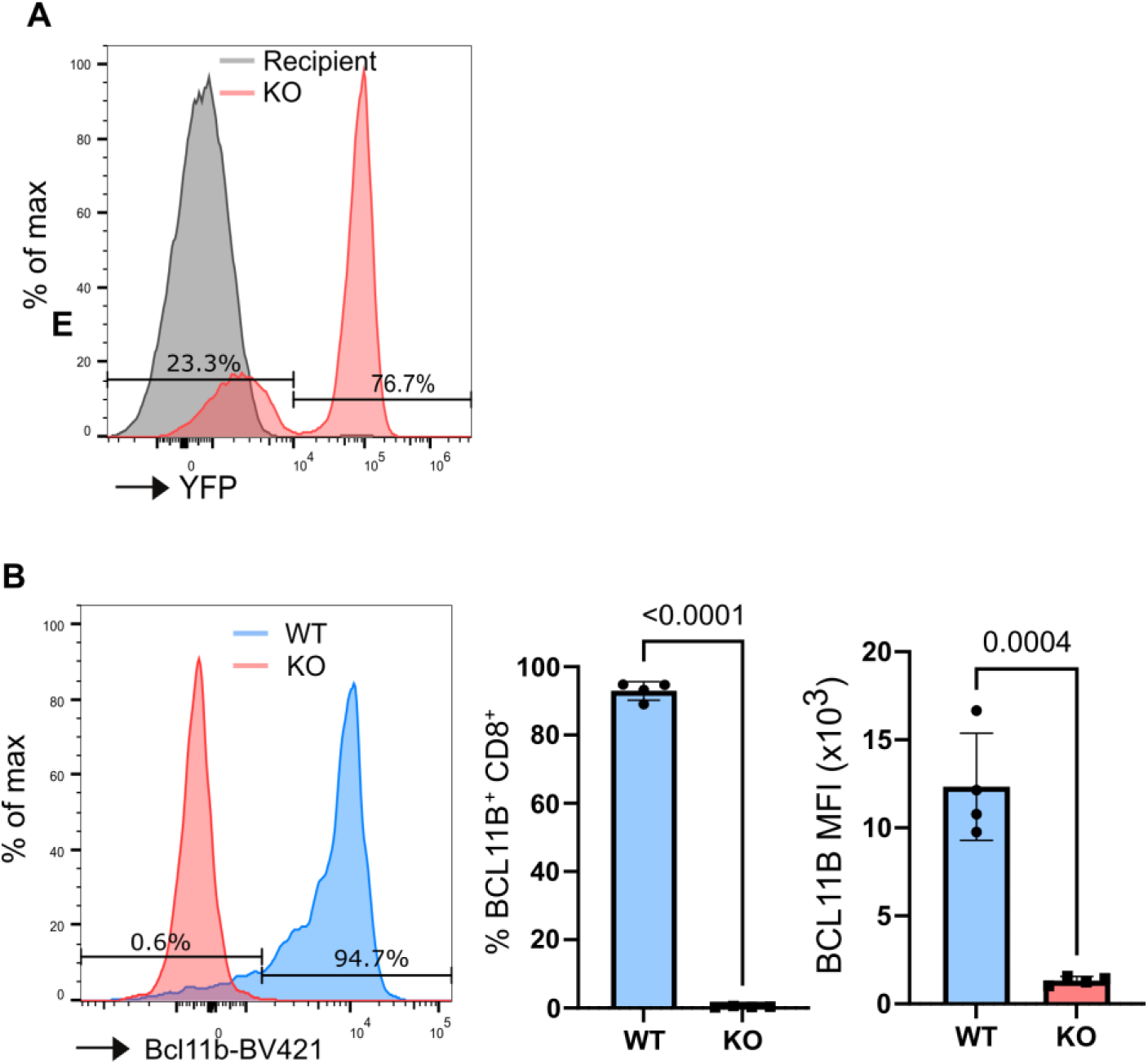
*Bcl11b* deletion with *GZMB*-Cre in TILs. **(A)** Histogram for YFP within the transferred CD45.2 *Bcl11b* KO OT-I CD8^+^ T cells versus recipient CD45.1 CD8^+^ T cells. **(B)** Histogram showing Bcl11b in transferred *Bcl11b* KO versus WT OT-I CD8^+^ T cells (left). Percentages and Bcl11b MFI quantification (central and right).

**Figure S3.**
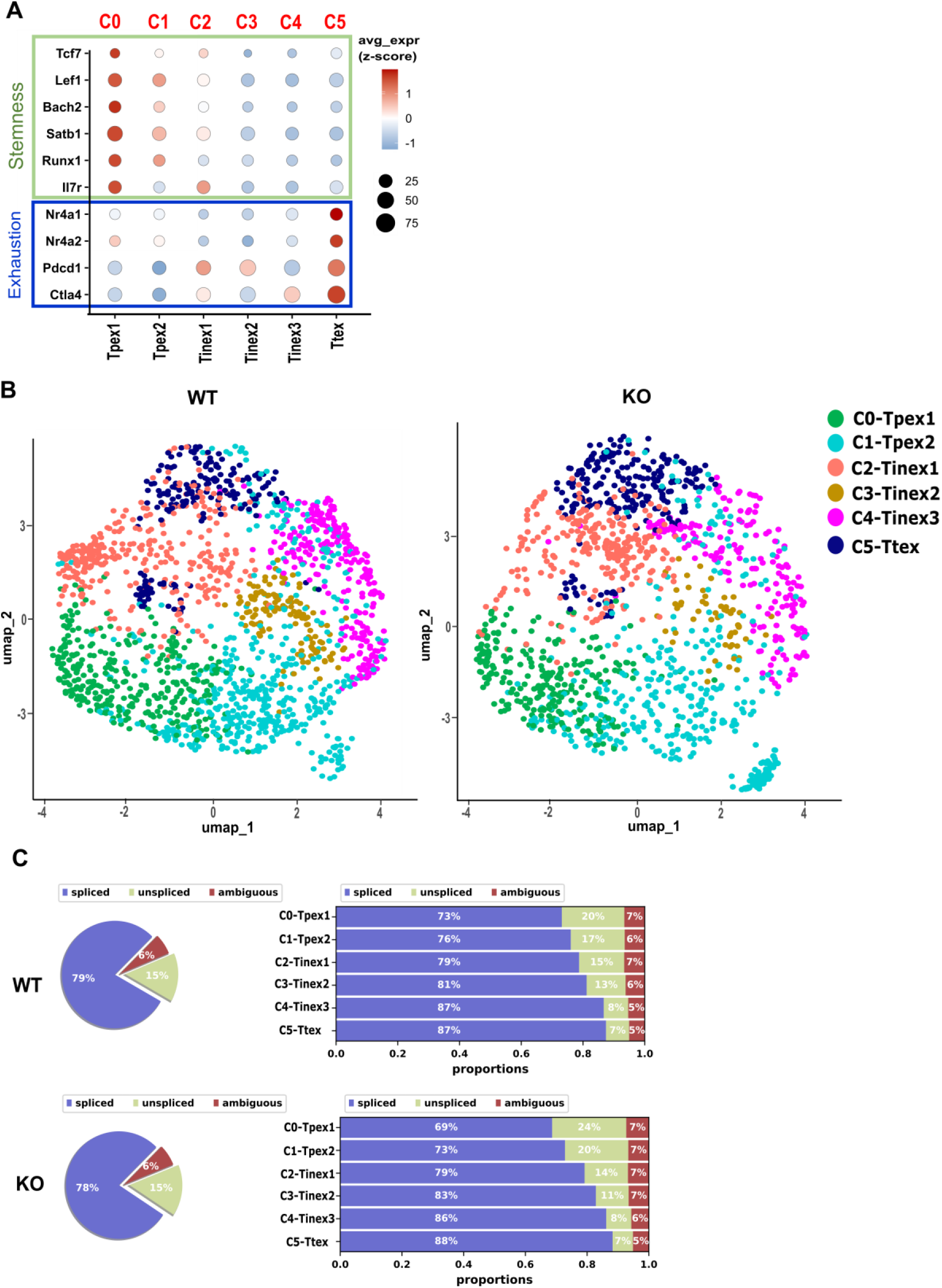
Cluster definition and splicing dynamics of transferred OT-I CD8^+^ TILs. scRNA-seq analysis of transferred OT-I CD8⁺ T cells isolated from peritoneal UPK10-OVA tumors. CD45.1 mice were injected i.p. with UPK10-OVA ovarian tumor cells and transferred with CD8^+^ T cells from donor CD45.2 OT-I *HGZMBCre*/*Bcl11b^f/f^*R26R-EYFP or *HGZMBCre*R26R-EYFP mice, activated as described in Fig. S1C. **(A)** Dot plot showing expression of definition marker genes for stemness and exhaustion across CD8⁺ TIL clusters. **(B)** UMAP projection and visualization of CD8^+^ T cells by graph-based cluster identity. Six transcriptionally distinct clusters, C0–C5, were identified and used for downstream marker-based annotation. **(C)** RNA velocity analysis illustrating predicted transcriptional dynamics, based on the relative abundance of spliced and unspliced transcripts, in *Bcl11b* KO and WT OT-I CD8⁺ TIL clusters.

**Figure S4.**
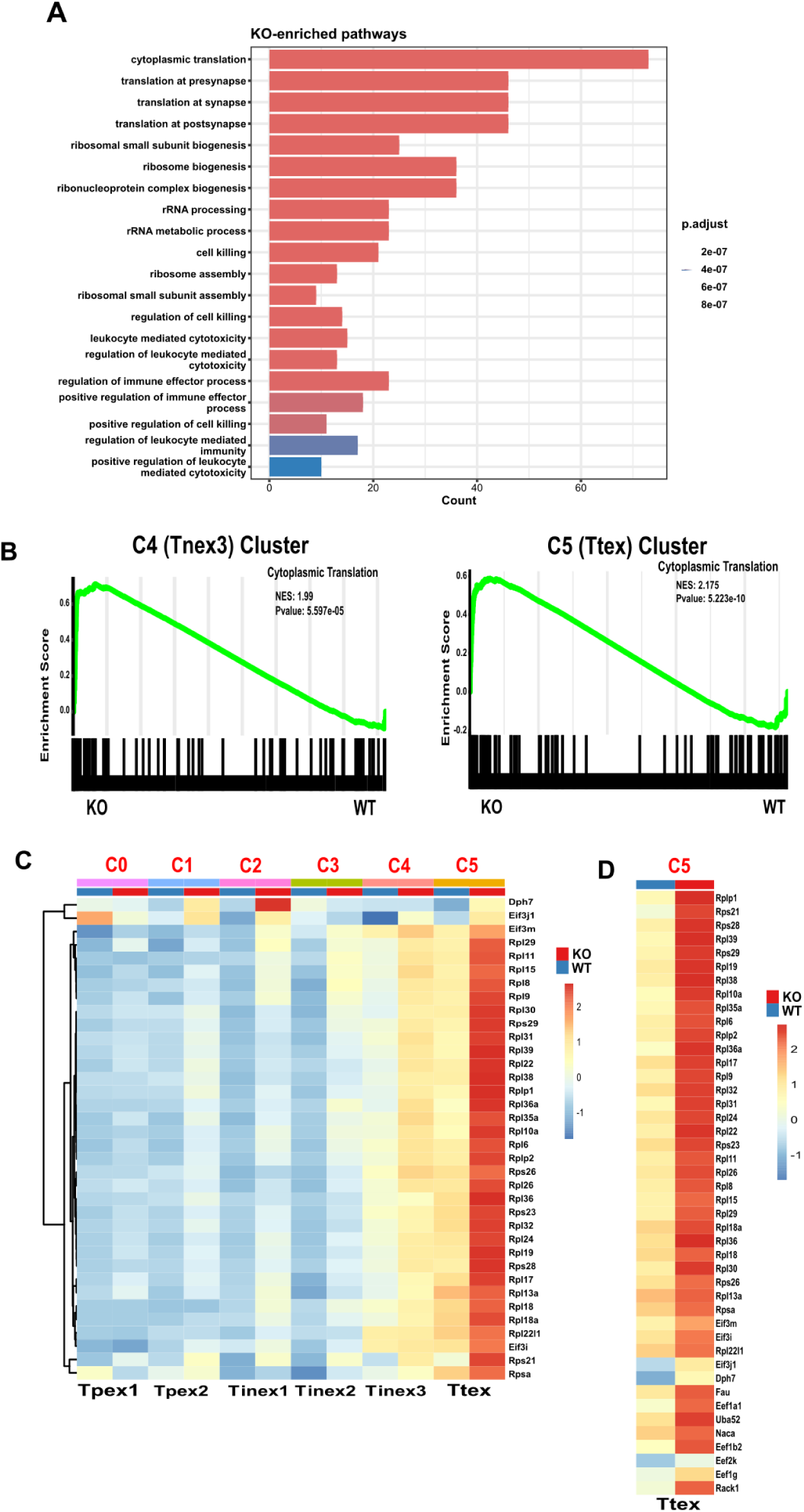
Genes associated with translation are enriched in *Bcl11b KO* C4 and C5 clusters. **(A)** Pathways enriched in *Bcl11b* KO clusters C4 and C5. **(B)** Gene set enrichment analysis (GSEA) of cytoplasmic translation pathway in *Bcl11b* KO and WT clusters C4 and C5 TILs. **(C)** Heatmap of genes associated with translation differentially expressed in *Bcl11b* KO versus WT CD8^+^ TIL clusters. **(D)** Heatmap of genes associated with translation differentially expressed in *Bcl11b* KO versus WT C5 CD8^+^ TILs.

**Figure S5.**
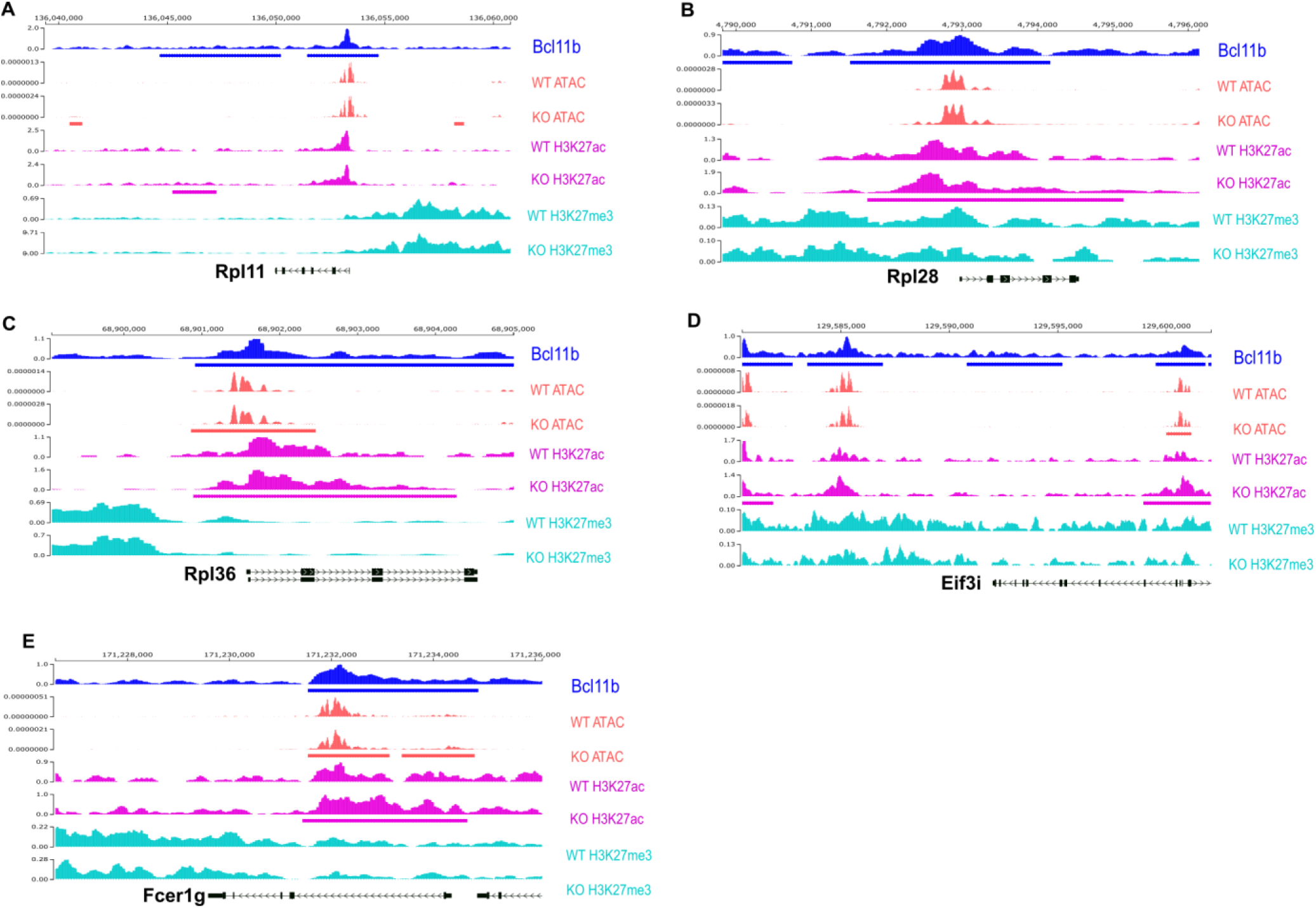
Bcl11b binding and epigenetic states at specific genes in CD8^+^ TILs. **(A-E)** Genomics tracks for ATAC-seq and CUT&RUN for Bcl11b, H3K27Ac and H3K27me3 at the indicated genes. Rectangles beneath each track indicate peaks that are significantly different between *Bcl11b* KO and WT TILs, or significantly bound by Bcl11b. CUT&RUN-seq for H3K27ac, H3K27me3, as well as ATACseq were conducted on polyclonal CD8^+^ TILs isolated from peritoneal UPK10-OVA tumors of *HGZMBCre*/*Bcl11b^f/f^*R26R-EYFP (KO) and *HGZMBCre*R26R-EYFP/CD45.2 (WT) mice. CUT&RUN-seq for Bcl11b was conducted on polyclonal CD8^+^ TILs isolated from peritoneal UPK10-OVA tumors of WT mice. Details are provided in Material and Methods, including for the analysis.

